# Estimating virulence from relative survival

**DOI:** 10.1101/530709

**Authors:** Philip Agnew

## Abstract

A pathogen’s virulence is a key parameter in the mathematical models on which most epidemiological theory is based. In these models virulence generally has a very specific definition where it is the increased per capita rate of mortality of infected hosts due to infection. Empirical studies involving the experimental infection of hosts often estimate virulence with the aim of comparing these estimates to values or patterns predicted in the theoretical literature. However most empirical studies do not estimate virulence as it is defined in the theoretical literature, thus potentially confounding comparisons between the two approaches. Here the analysis of relative survival is applied to the type of data routinely generated in empirical studies to estimate virulence as it is defined in the theoretical literature. The theoretical grounds for approach are outlined, followed by worked examples estimating the virulence of different pathogens with data from published studies. Code allowing virulence to be estimated by maximum likelihood with *R* is provided.

## Introduction

Empirical studies involving the experimental infection of hosts often estimate the pathogen’s virulence and compare these estimates to values or patterns predicted in the theoretical literature. However most empirical studies do not estimate virulence as it is defined in the mathematical models on which most epidemiological theory is based [1–3]. Instead alternative or proxy measures of virulence are used and they vary in how well they correlate with virulence as it is defined in the theoretical literature [4, 5]. Consequently it is questionable as to how valid most comparisons between data and theory are where virulence is concerned [4].

Here the analysis of relative survival is presented as a means to estimate pathogen virulence, as it is generally defined in the theoretical literature, from the type of data routinely generated in empirical studies recording the survival of experimentally-infected hosts over time.

### Empirical context

Empirical studies investigating the effects of pathogens on hosts often involve the initial establishment of several uninfected populations of hosts, experimentally-infecting some of them, and recording how host survival changes over time in the different treatments. Statistical analyses are then applied to compare survival in the infected vs. uninfected treatments.

The members of the initial uninfected populations are usually chosen such that the expected pattern of mortality in the different populations would be expected to be the same were it not for the experimental intervention of infection. Thus, all else being equal, any reduction in survival for infected populations can be attributed to the effect of infection.

The pattern of events occurring in this type of experimental design can be described by the same expressions as those used in population dynamics models describing the flow of hosts between different compartments, e.g., susceptible-infectious-recovered (SIR) models. The dynamics described by such models can be complex and open to influence by many parameters. However in the type of experiment outlined above it is often possible to simplify matters such that the only population dynamics to be recorded are those for how host population sizes change over time due to host mortality. For example, the pattern of events in an uninfected (*X*) and infected (*Y*) population of hosts can be expressed as,

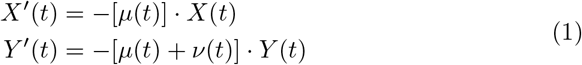

where *X′*(*t*) and *Y′*(*t*) are the rates of change in the size of the uninfected and infected populations at time *t*, respectively. These are both negative and determined by the size of each population at time *t*, *X*(*t*) and *Y*(*t*), respectively, multiplied by the average rate at which hosts die. In the uninfected population this rate is determined by the ‘natural’ or background rate of mortality at time *t*, *μ*(*t*). In the infected population there is an additional rate of mortality, *ν*(*t*), which is the average increase in the rate of mortality of infected hosts due to infection at time t; this is how virulence is defined in most epidemiological models on which the theoretical literature is based [6, 7].

Here the population dynamics have been simplified by assuming all hosts are of the same age, where a host’s age and that of their infections is set to zero at time to, when survival is first recorded; hence age and time are interchangeable in the following expressions. It is also assumed there are are no births during the experiment and the observed dynamics are not influenced by any density-dependent processes.

There are no dynamics describing the gain of infection, as it is assumed an individual’s infection status was determined in the period prior to survival being recorded and there is no transmission of infection during the experiment, e.g., because infected and uninfected hosts are housed separately. Unless stated otherwise, these models assume all hosts in an infected treatment are infected.

There are also no dynamics describing the loss of, or recovery from, infection. This is an influential parameter for the predictions of many epidemiological models. The initial models below assume hosts cannot recover from infection or do not do so within the time-frame of the experiment. Subsequent models allow for recovery from infection and other sources of variation influencing the observed pattern of mortality in an infected treatment or population. Allowing for such variation can qualitatively change the estimated pattern of virulence experienced by individual hosts relative to the pattern observed for an infected population or treatment as a whole.

### Survival

An alternative means of describing the population dynamics within an experiment is with survival functions where time is continuous. For example, the cumulative survival function *S(t)* is the probability and individual in a particular population at the beginning of an experiment, *t*_0_, will still be alive at time *t*,

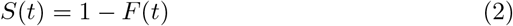

and is the complement of the cumulative density function, *F(t)*, for the probability an individual will have died by time *t*; 0 ≤ *S*(*t*), *F*(*t*) ≤ 1.

Differentiating *F(t)* with respect to time gives the rate at which mortality reduces the size of the population at time *t*,

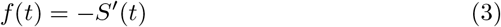

where *f(t)* is the probability density function for mortality. It corresponds with the number of individuals dying at time *t*, divided by the initial size of the population. The number of individuals dying at time *t* is generally the data collected in this type of experiment.

Whereas *f(t)* represents the probability an individual alive at *t*_0_ will die at time *t*, the hazard function, *h(t)*, represents the probability an individual alive at time *t* will die at time *t*,

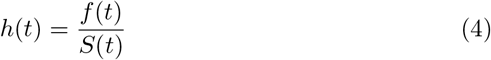

where the probability of dying at time *t*, *f(t)*, is corrected by the probability of being alive at time *t*, *S(t)*. This is the rate of mortality in the population at time *t* and represents the per capita risk of dying at time *t*.

If the hazard function *h*(*t*) represents the risk an individual alive at time *t* will die at time *t*, the cumulative hazard function, *H* (*t*), represents the individual’s accumulated exposure to the risk of dying at time *t*. It is related to the cumulative survival function, *S* (*t*), as,

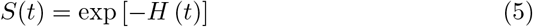

and can take values greater than one.

Analyses testing for the effects of a pathogen on the survival of infected vs. uninfected hosts usually involve the estimation and comparison of one of the expressions above. What makes the analysis of relative survival different is how it treats the survival of infected hosts.

### Relative survival

The analysis of relative survival is frequently encountered in the medical literature [8–11], where it is the method of choice for estimating how survival in populations of patients is affected by a particular disease or illness, e.g., different types of cancer [12, 13]. What the relative survival approach does is to assume individuals in the target population are exposed to two independent and mutually exclusive sources of mortality;

i. *Background or ‘natural’ mortality*. This is the mortality individuals in the target population would be expected to experience had they not been afflicted by the disease or illness in question, and,
ii. *Mortality due to disease or illness*. This is mortality individuals in the target population experience due to the disease or illness in question.

The following outlines the relative survival approach applied to describing survival in infected vs. uninfected populations of hosts.

When infected hosts die, it is not possible to tell whether they died due to background mortality or due to infection. However to remain alive means the host has not died due to the cumulative effects of background mortality, *F_BCK_* (*t*), or the cumulative effects of mortality due to infection, *F_INF_* (*t*). As these two sources of mortality are independent, the probability an infected host will be observed surviving until time *t*, *S_OBS.INF_* (*t*), can be calculated as,

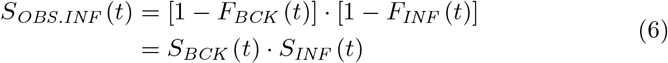

where *S_Bck_* (*t*) and *S_INF_* (*t*) are the cumulative survival functions for background mortality and mortality due to infection at time *t*, respectively.

The relative survival of infected hosts at time *t*, *S_REL_* (*t*), is calculated as their observed probability of surviving until time *t*, divided by that for uninfected hosts,

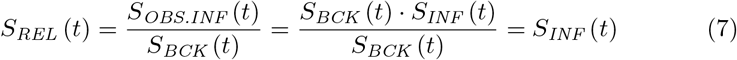

which equals the expected survival of infected hosts due only to the effects of infection, *S_INF_* (*t*). That is, relative survival is the observed survival of infected hosts corrected for background mortality.

If there is no background mortality, as happens in some experiments, *S_BCK_* (*t*) = 1 and the observed mortality in an infected treatment can be analysed directly as being due to the effect of infection; *S_REL_* (*t*) = *S_OBS.INF_* (*t*) = *S_INF_* (*t*) (7).

Differentiating *S_OBS.INF_* (*t*) with respect to time and taking the negative gives the probability density function for mortality observed in the infected treatment at time *t*, *f_OBS.INF_* (*t*),

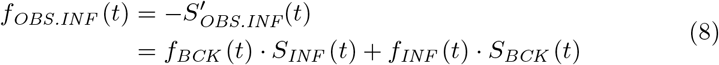

where *f_BCK_* (*t*) and *f_INF_* (*t*) are the probability density functions for the probability an individual alive at time *t*_0_ will die at time *t* due to background mortality or mortality due to infection, respectively.

Dividing *f_OBS.INF_* (*t*) by *S_OBS.INF_* (*t*) gives the hazard function, *h_OSS.INF_* (*t*),

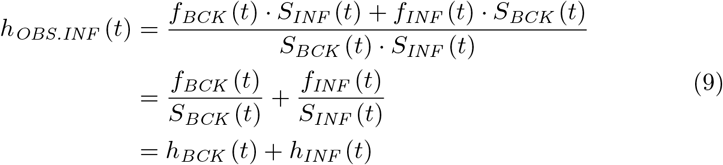

showing the observed rate of mortality in the infected population is the sum of the background rate of mortality at time *t*, *h_BCK_* (*t*), plus the rate of mortality due to infection at time *t*, *h_INF_* (*t*), that is, the pathogen’s virulence at time *t*.

Re-arranging (1) for infected hosts shows the population dynamics describing the per capita rate of decrease in the size of the infected population, −*Y′*(*t*)/*Y* (*t*), equals the relative survival expression for the rate of mortality observed in the infected population (9),

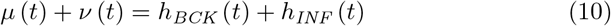

The following sections illustrate how these rates can be estimated by analysing the relative survival of infected and uninfected hosts.

## Estimating virulence

### Estimates from survival curves

Published figures showing survival curves for infected and uninfected populations of hosts can be used to estimate a pathogen’s virulence. The observed survival in an infected population at time *t*, divided by that in a matching uninfected population estimates the relative survival of infected hosts at time *t*, *S_REL_* (*t*). Differentiating *S_REL_* (*t*) with respect to time and taking the negative gives the probability density function for the probability an infected host alive at time *t*_0_ will die due to infection at time *t*, *f_INF_* (*t*),

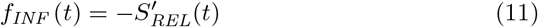

Dividing *f_INF_* (*t*) by the relative survival in the infected population at time *t*, *S_REL_* (*t*), gives the hazard function, *h_INF_* (*t*), at time *t*,

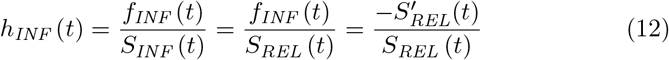

thus a pathogen’s virulence can be estimated directly from survival curves by using them to calculate the relative survival of infected hosts at time *t* and the rate of change in relative survival at time *t*.

This approach to estimating virulence does not require any information on the frequency of infected or uninfected individuals dying or remaining alive at any point in time. However, without information on the actual frequencies of individuals dying or remaining alive at different points in time, it is not possible to calculate the variance associated with these estimates of virulence. Consequently it is not possible to tell if, or when, these estimates differ from one another, or from zero.

### Estimates at individual points in time

When sampling is carried out at regular intervals, the rate of mortality in interval *i*, *h*(*i*), can be estimated as,

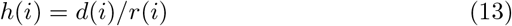

where *d*(*i*) is the number of individuals dying during interval *i* and *r*(*i*) is the number of individuals alive, or at risk of dying, at the beginning of interval *i*. Re-arranging (9) shows the difference in rates of mortality observed for infected and uninfected treatments at a given time estimates the rate of mortality due to infection at the time, hence these differences for the interval *i*;

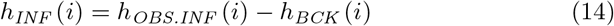

estimate the pathogen’s virulence in the interval.

These estimates of virulence are based on the difference between two binomial proportions. There are many methods to calculate the confidence intervals for this type of difference. Wald’s confidence interval is the most widely used, but it is not recommended by some authors [14]. An alternative is to calculate Wald’s modified or adjusted confidence intervals as proposed by Agresti & Caffo [15]; see supplementary file S01 for details.

If there is no mortality recorded in a control population the observed mortality of infected hosts can be directly analysed as being due to infection. Poisson regression models may provide a useful means for estimating a pathogen’s virulence in individual intervals as they can make use of functions already available in statistical software packages, including the influence of random effects (e.g., see Austin [16]).

There are limitations to estimating virulence at individual points in time. One is that generating a series of individual estimates based on small subsets of a larger dataset is not an efficient means of exploiting the available data. Another is differences among individual estimates of virulence are potentially confounded with random variation in mortality rates over time. Such variation can influence estimates towards the beginning of an experiment when mortality rates are low in both infected and uninfected treatments. However it is likely to be more influential as the number of individuals within a treatment decreases and stochastic variation in the number of individuals dying has more influence on the proportion of individuals dying (or remaining alive) within a sampling interval.

An alternative to estimating virulence at individual points in time is to estimate it as a function of time.

### Estimates as a function of time

Probability distributions can be used to describe the probability an event will occur between two points in time. When the event is death, these distributions allow survival, probability density, and hazard functions to be described as continuous functions of time.

Cumulative survival functions described by different probability distribution all decrease monotonically over time from 1 to 0. However these functions are specified differently for each probability distribution, as are the corresponding probability density and hazard functions. This variation allows different probability distributions to describe different patterns of survival over time. Furthermore the functions for each probability distribution require at least one parameter to be specified and varying the values of these parameters provides flexibility in the pattern of survival each probability distribution can describe.

Figure 1 illustrates some of the variation possible in the patterns of survival and mortality of different probability distributions. Here the survival, probability density and hazard functions are plotted for the exponential, Weibull, Gumbel and Frechet distributions when survival was constrained such that 5% of the original population remained alive on day 14; *S*(*t_14_*) = 0.05. Given this constraint, survival curves varied from being concave to convex (Fig 1 a-d). The probability density function that an individual alive a time *t*_0_ would die at time *t* varied from decreasing monotonically to having a unimodal distribution skewed to either the left or right (Fig 1e-h). The corresponding hazard functions varied from being constant (Fig 1i), to increasing at a constant rate (Fig 1j), increasing at an accelerating rate (Fig 1k), or having a unimodal pattern increasing and then decreasing over time (Fig 1l).

**Figure 1:**
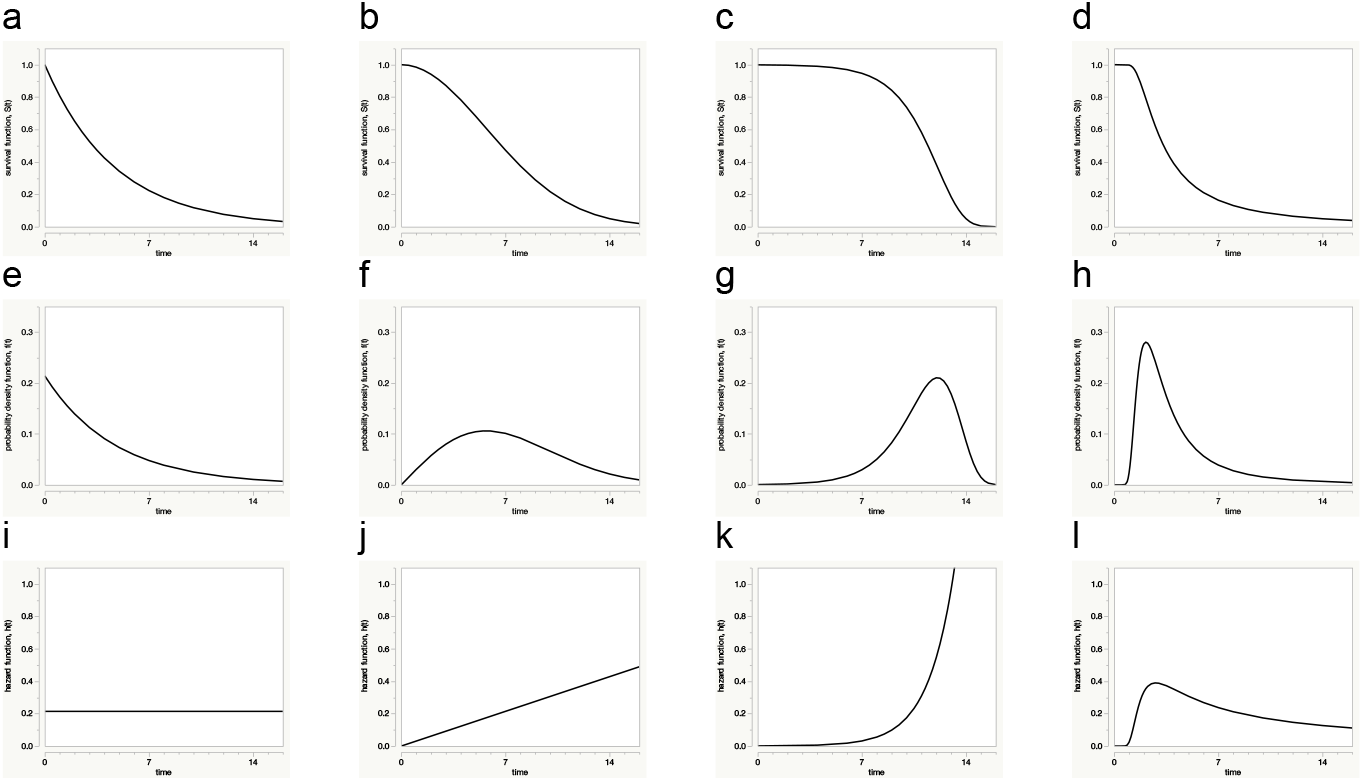
Survival (a-d), probability density (e-h) and hazard (i-l) functions for the exponential (column 1), Weibull (column 2), Gumbel (column 3) and Frechet (column 4) distributions. Note the scale of Y-axis for the probability distribution figures (e-h) is different to the scale used for the survival and hazard functions.

Table 1 gives the survival, probability density and hazard functions corresponding with each of the four probability distributions above. There are different ways these functions can be expressed. Here they are expressed in terms of location (*a*) and scale (*b*) parameters which determine where and over what range the data are distributed along the time axis, respectively; the exponential distribution is a special case of the Weibull distribution when *b* =1.

Values of *a* and *b* are interchangeable between functions, such that, estimates of *a* and *b* from cumulative survival data can be plugged into the hazard function for the estimated of the rate of mortality over time, and vice-versa.

**Table 1:**
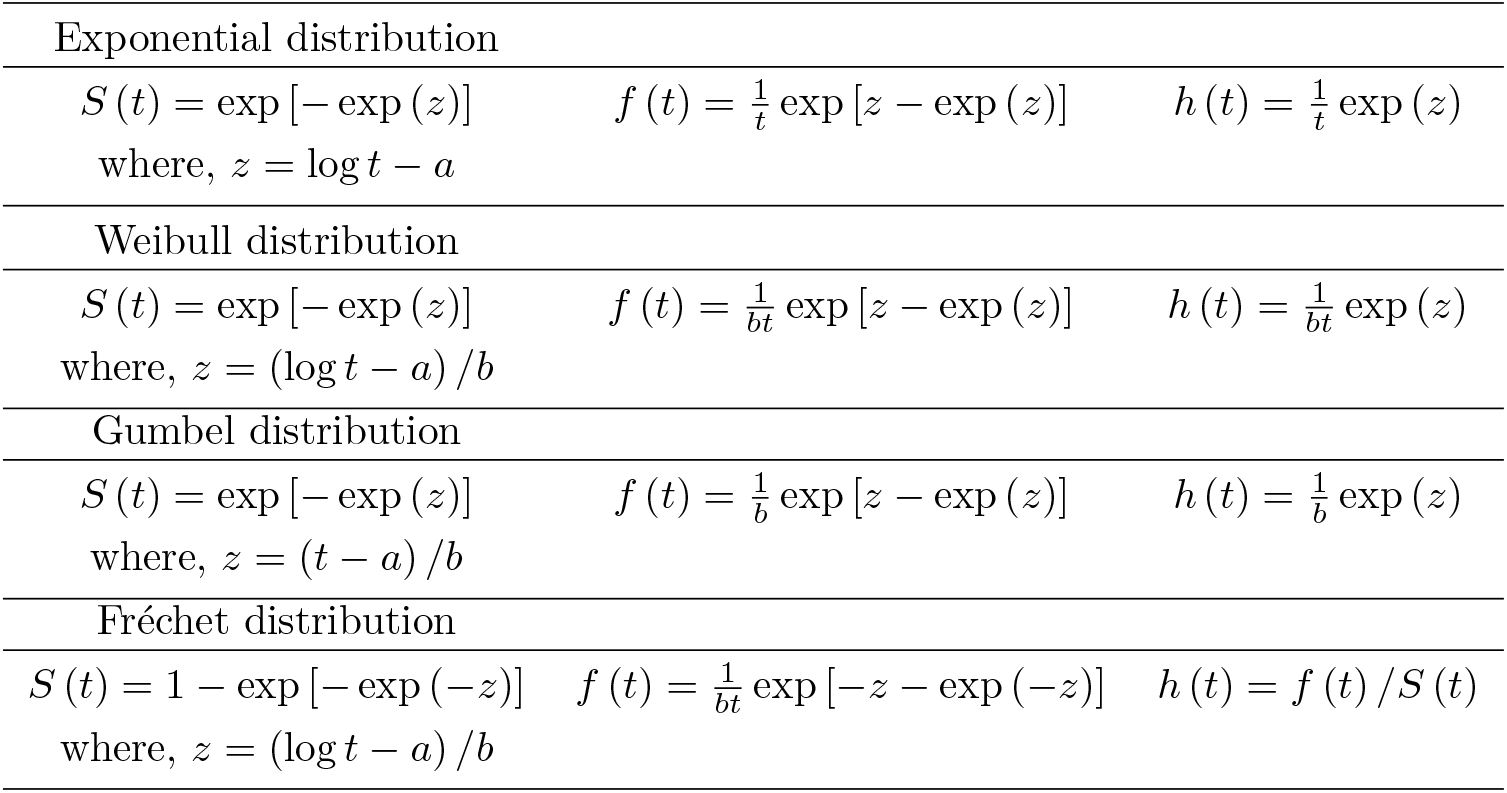
Survival, probability density and hazard functions for the Exponential, Weibull, Gumbel and Frechet distributions

The value of these parameters can be made to depend on covariates, such as, dose, temperature, gender, etc…

Various approaches can be used to determine the parameter values giving the closest fit between survival functions and survival data. The next section outlines the approach using likelihood models.

#### Likelihood models

Data collected during the course of a survival experiment provides information on the frequency of individuals dying between sampling intervals. These can be used to estimate the probability density function, *f(t)*, for the probability an individual alive at time t0 will die at time *t*. However empirical studies are frequently terminated before all, or even most, individuals die. Consequently these individuals do not experience the event of interest. Nonetheless it is known they survived at least until the end of the experiment and this information can contribute towards estimates for the probability of dying at time *t*. These are known as censored individuals, or more precisely right-censored individuals. They include individuals removed from populations during the course of an experiment, e.g. to control for infection success, those that escape, or are accidently killed, where the timing of the event is known.

If the probability of dying at any one time *t* is equal for all *n* members of a population and the death of any one individual *i* has no effect on the timing of death of any other individuals, the overall likelihood *L* of dying at time *t* is the product of the probability of each individual dying at time *t*,

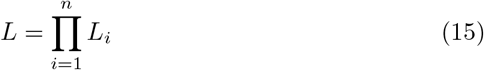

where *L*_*i*_ = *f* (*t_i_*). This allows for right-censored individuals when expressed as,

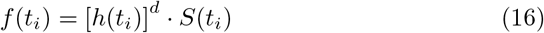

with *d* being a death indicator taking a value of 1 for individuals dying during the experiment and 0 for those censored during or at the end of the experiment.

It is often more convenient to work with the likelihood expression after log-transformation,

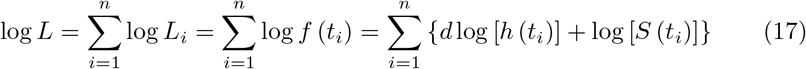

this log-likelihood expression can be used for analysing survival data, allowing for right-censored individuals. Substituting the survival and hazard functions in (17) with equations (6) and (9), respectively, gives

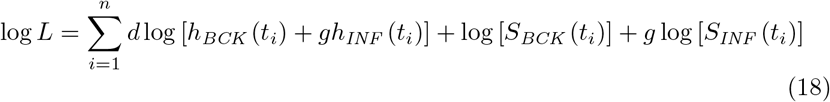

where *g* is an indicator of infection taking a value of 1 for individuals exposed to infection and 0 for those not exposed to infection. This log-likelihood expression can be used for analysing relative survival, allowing for right-censored individuals.

Approximate confidence intervals for the hazard function describing the pathogen’s virulence can be estimated using the delta method [17]; see S02 for details.

The goodness of fit between data and the likelihood model describing them is reflected in the loss function; the closer to zero the loss, the better the fit. Likelihood-ratio tests can be used to test for significant differences in the loss of nested models where one model needs to be a simplified version of the other, e.g., models with or without interaction terms. Alternatively models can be compared based on criteria determining which of the rival models is best, e.g., Akaike’s information criterion (AIC). An advantage of these approaches is they allow non-nested models to be compared, e.g., when the same data are analysed with models based on different probability distributions. Here Akaike’s information criterion corrected for finite sample sizes, AICc [18], is used to compare models.

Supplementary file S03 gives details of how to specify (18) in *R* [19] and estimate parameter values by maximum likelihood using the *bbmle* package by Ben Bolker and the R Core development team [20]. Supplementary file S04 shows how to specify (18) for analysis with the non-linear platform of JMP [21].

Maximum likelihood estimation techniques used to solve likelihood problems require initial values for the parameters to be estimated. Some probability distributions lend themselves to this task by having functions that can be transformed into linear functions of time. This allows initial parameter values to be estimated by ordinary linear regression; see S05.

## Worked examples

This section presents the results of analyses where virulence has been estimated from the data of published studies. The aim is to illustrate the analysis of relative survival rather than to provide definitive analyses of the data or to challenge the original analyses. Supplementary files provide details of each analysis.

### Virulence increasing over time

Here the rate of mortality due to infection is estimated as increasing over time at an accelerating rate.

Blanford et al. [22] exposed replicate populations of adult female *Anopheles stephensi* mosquitoes to a total of 17 different isolates of four fungal pathogens and recorded their survival over the next 14 days. Simon Blanford kindly provided the original data from the experiment and Matthew Thomas generously allowed their inclusion as supplementary material in this study. Here a subset of the data is analysed where the survival of adult female mosquitoes exposed to isolate *Bb06* of *Beauveria bassiana* is compared to that of uninfected females in matching control treatments.

Most of the females exposed to infection died within the 14 days of the experiment, whereas roughly half of those in the uninfected cages died (Fig 2a). Consequently roughly half of the mortality observed in the infected treatment was expected to be due to background mortality (7).

**Figure 2:**
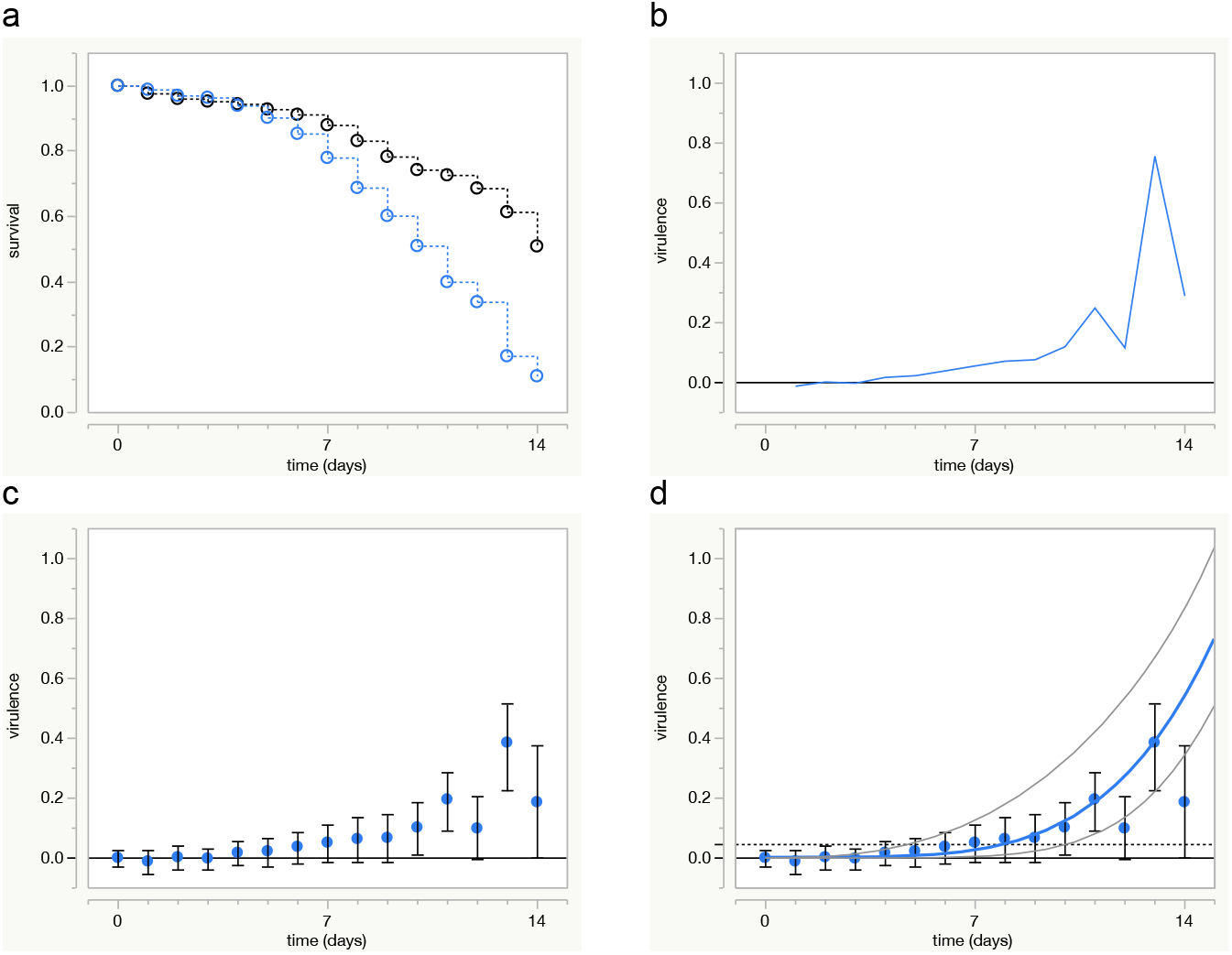
Survival and estimates of virulence for mosquitoes with fungal infections. (a) Survival of uninfected (black) and infected (blue) mosquitoes pooled across replicate cages, (b) Virulence as estimated from dynamics of relative survival, (c) Virulence estimated at individual points in time (±95% *c.i*.) (d) Maximum likelihood estimate of virulence (blue line ±95% c.i. grey lines). The dotted horizontal line is the estimate for virulence assuming constant rates of mortality. Symbols as in (c).

The pathogen’s virulence was estimated as tending to increase over time based on the rate of change in relative survival calculated from survival curve data (12) (Fig 2b). Estimates based on the proportion of individuals dying in each daily interval (13) confirmed this pattern and estimated virulence as being significantly greater than zero in the second week of the experiment, although not consistently so (Fig 2c).

Initial investigations of the data suggested the Weibull distribution was appropriate for describing background mortality and mortality due to infection, thus the log-likelihood expression for analysing relative survival (18) was parameterised accordingly; see S06. In numerical terms the hazard function estimating the pathogen’s virulence was,

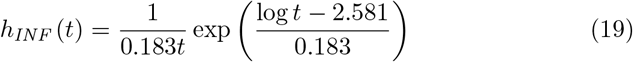

As, 0 < (*b_INF_* = 0.183) < 0.5, this Weibull hazard function describes the rate of mortality due to infection as increasing monotonically over time at an accelerating rate (Fig 2d).

In contrast to estimates at individual points in time, the delta method estimated the lower 95% confidence interval as consistently greater than zero in the second week of the experiment (Fig 2d).

#### Comparison with a constant rate of mortality model

Epidemiological models commonly assume mortality rates remain constant over time. This assumption is sometimes justified by observed data, but it is more frequently a convenience making models easier to manipulate and interpret. For example, a pathogen’s basic reproduction number *R*_0_ is often calculated as being proportional to the average longevity of infected hosts [6, 7]. This can be calculated from the area under a survival curve when there are no censored individuals or by integration of the survival function when there are; see S07. When mortality rates are constant, the calculation simplifies to, 1/(*μ* + *ν*), or, 1/(*h_BCK_* + *h_INF_*).

The likelihood model above was re-run with *b_BCK_* = *b_INF_* = 1; thus constraining the Weibull hazard functions to constant rates of mortality. The constrained model estimated the background rate of mortality as 0.041 day^-1^ and that due to infection as, 0.047 day^-1^ (Fig 2d). The corresponding estimates for the average longevity of infected hosts were, 1/(0.041 + 0.047) ~ 11.4 days for the constrained model and ~ 10.3 days for the unconstrained model; see S06. All else being equal, estimates of *R_0_* based on the assumption of constant mortality rates would be ~ 10% greater than those based on more realistic estimates of host mortality.

### Proportional virulence

Here a pathogen’s virulence is estimated as proportional to the dose of infection hosts were exposed to.

Proportional hazards models are a popular means of comparing survival in infected vs. uninfected populations of hosts. An attractive feature of this approach is to condense differences in host mortality into a single and easy to interpret value, e.g., infection doubled a host’s risk of dying. These models assume there is a underlying rate of mortality experienced by all individuals under study, with the difference in the mortality rates among treatments or populations being determined by their deviation from this underlying rate. Non- or semi-parametric proportional hazards models make no assumptions as to the distribution of the underlying rate, only quantifying the deviation from it. In parametric models the underlying rate of mortality is specified according to a particular probability distribution. In both cases the hazard functions for the populations under study need to satisfy the relationship,

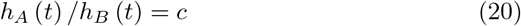

where *A* and *B* are independent populations and *c* is a constant, i.e., the ratio of mortality rates in the populations being compared remains constant over time [23].

The data in this example are from a study by Lorenz & Koella [24] and are freely available [25]. Larvae of the mosquito *An. gambiae* were exposed to spores of the microsporidian parasite *Vavraia culicis* in six different dose treatments, and there was an unexposed control treatment. Larvae were reared in individual vials on diets of high or low food availability. As adults they remained in their vials of origin and were provided with sugar-water. Adult longevity was recorded daily until all individuals died. The relative survival of adult females in all dose treatments is analysed, however for clarity, only data from the control, lowest and highest dose treatments are presented in the accompanying figure.

Larval food availability had little effect on adult longevity, whereas infection reduced survival and tended to do in dose-dependent manner (Fig 3a). Two females in the uninfected, high larval food availability treatment died within three days of emerging. This skewed the lower tail of the probability density function for background mortality towards a pattern described by the Gumbel distribution (c.f. Fig 1g). The Weibull distribution was used to describe mortality due to infection. Different models were fitted to the data, with the results of the best model according to AICc presented below; see S08.

**Figure 3:**
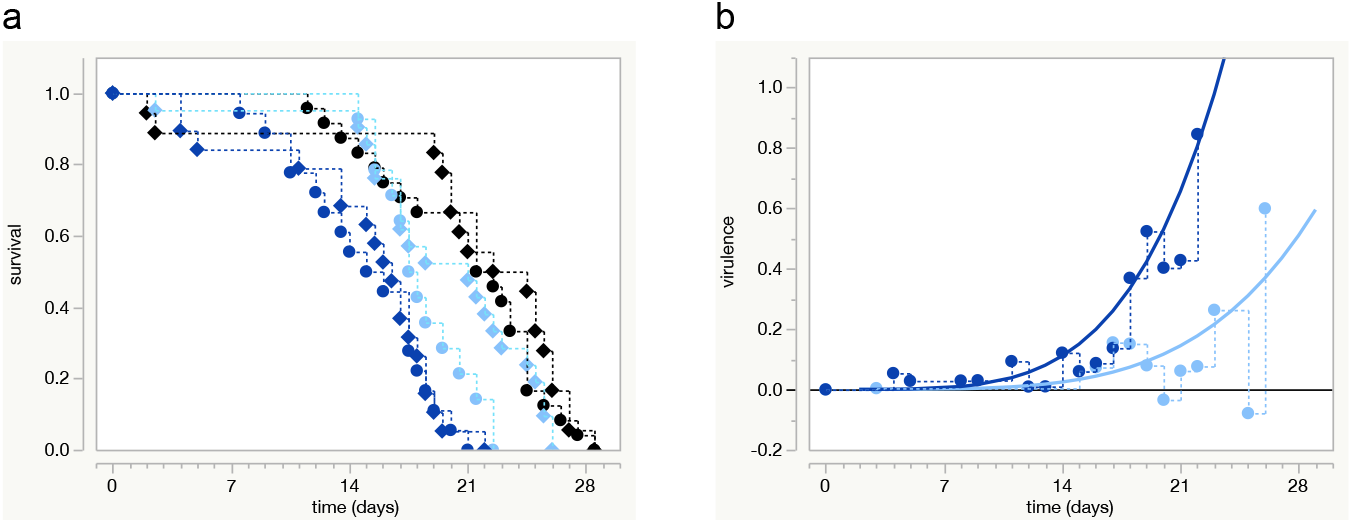
Proportional virulence. (a) Survival in control (black), lowest dose (light blue) and highest dose (dark blue) treatments. Low and high food treatments in circles and diamonds, respectively, (b) Observed and estimated virulence in the lowest and highest dose treatments. Observed data (stepped lines, symbols) based on daily data pooled for larval food treatments. Smooth curves show maximum likelihood estimates for virulence.

The best model pooled data from the two larval food treatments and estimated the rate of mortality due to infection as a log-linear function of the dose of spores larvae were exposed to;

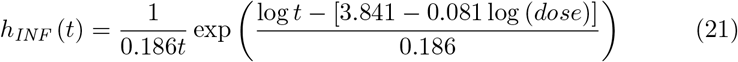

As in the previous example, the scale parameter estimated for the Weibull hazard function was, 0 < (*b_INF_* = 0.186) < 0.5, meaning rates of mortality due to infection were estimated as increasing monotonically over time at an accelerating rate (Fig 3b). The estimate for the location parameter, *a_INF_* = 3.841 − 0.081 log (dose), has the effect of bringing forward the scheduled pattern of mortality as dose increases.

The pathogen’s virulence was also estimated as being proportional to the log (dose) of infection: Weibull hazard functions are proportional when their scale parameters are equal as their ratio reduces to,

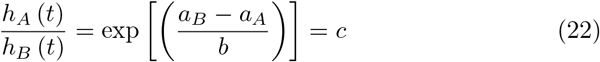

where *A* and *B* are independent populations, *b_A_* = *b_B_* = *b* and *c* a constant.

This was not the case for the observed rates of mortality in the infected vs. uninfected treatments. However separating out the contribution of background mortality to the observed mortality of infected hosts resulted in a single value of *b_INF_* being estimated for all six dose treatments. Consequently the relative survival approach estimated the rate of mortality due to infection as being proportional to the dose of infection. In numerical terms, the pathogen’s virulence was estimated as increasing by ~ 4.5 times between the lowest and highest dose treatments (Fig 3b).

Supplementary file S08 illustrates how the choice of different probability distributions influences the goodness of fit between the log-likelihood model and the observed data.

Supplementary file S09 illustrates how the data above can be considered in terms of an accelerated failure time (AFT) model where dose acts to scale the passage of time for mortality due to infection.

### Unimodal virulence

This example estimates the virulence for three isolates of a fungal pathogen as increasing and then decreasing over time.

The data are from the study by Blanford et al. [22] and involve the fungal pathogen *Metarhizium anisopliae*. Isolates *Ma06, Ma07* and *Ma08* were each used to infect replicate host populations and there were replicate control populations unexposed to infection in the same block of the experiment; see S10.

There was less background mortality in this block of the experiment (Fig 4a) than that analysed above (c.f. Fig 2a). Nonetheless background mortality will have influenced the observed mortality of mosquitoes in the infected treatments.

**Figure 4:**
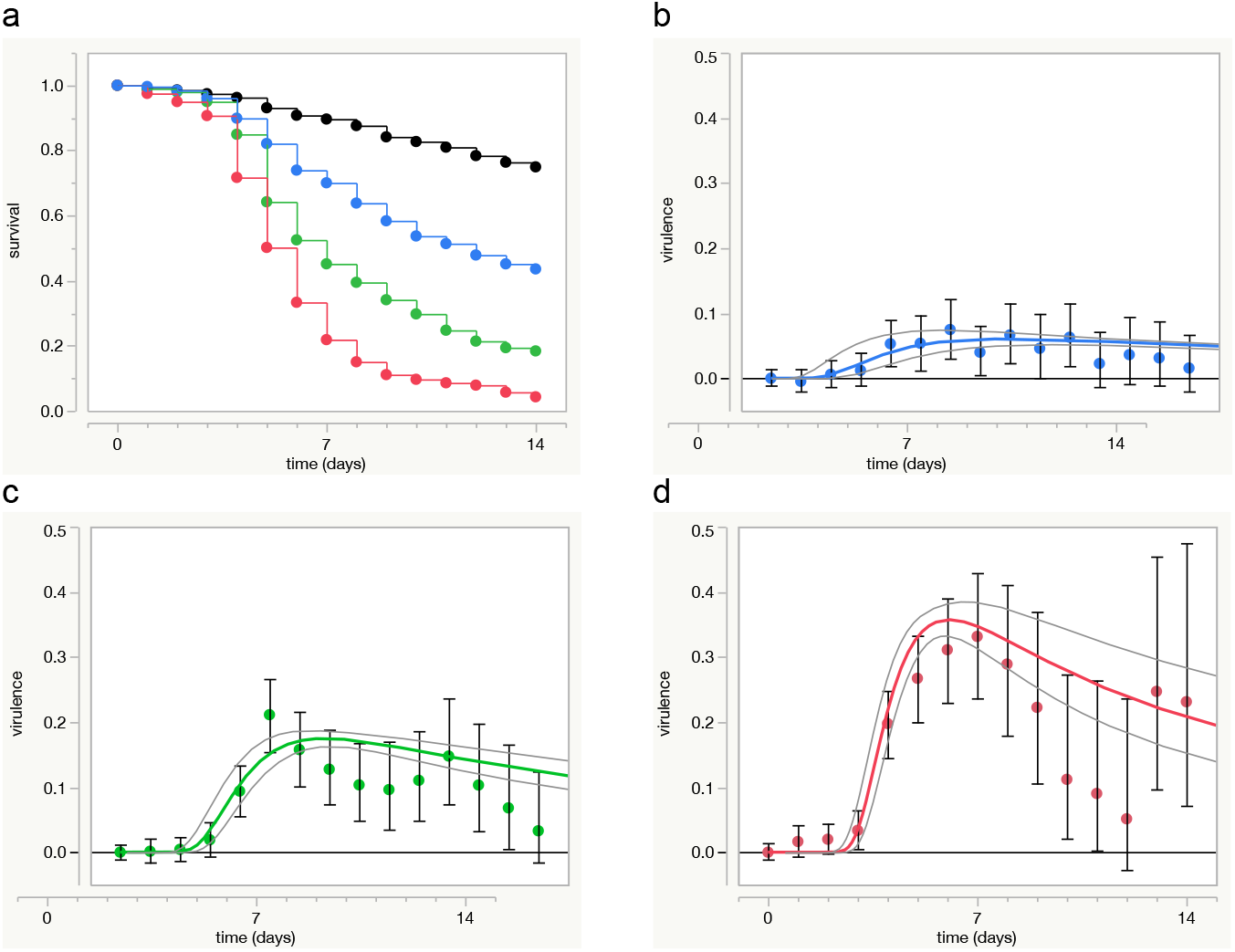
Unimodal patterns of virulence for three fungal isolates. (a) Survival in the control treatment (black), and those exposed to isolates *Ma06* (green), *Ma07* (red), *Ma08* (blue), (b-d) Daily estimates of virulence for each isolate (symbols ±95% c.i.) and maximum likelihood estimates (coloured line ±95% c.i. grey lines); colours as in (a).

The Weibull distribution was considered suitable for describing the background mortality. However survival curves tended to flatten out over time in the infected treatments, particularly for isolate *Ma07* (Fig 4a). This flattening corresponds with the rate of mortality decreasing over time. Daily estimates for the virulence of each isolate agreed with this pattern as they initially increased and then tended back towards zero (Fig 4b-d). This is not a pattern of mortality the Weibull or Gumbel distributions can describe, but one the Frechet distribution can (c.f. Fig. 1l).

The log-likelihood expression (18) was parameterised where the location and scale parameters for the background mortality, *a_BCk_* and *b_BCK_* respectively, were estimated using Weibull distribution functions. Those for mortality due to infection, *a_INF_* and *b*_INF_, were estimated separately for each isolate with Frechet distribution functions.

Estimates of *a_INF_* and *b_INF_* for each fungal isolate (±95% *c.i.)* were nonoverlapping and chosen as the best values for each isolate; see S10. Each described a unimodal pattern of virulence increasing and then decreasing over time for each isolate (Fig 4b-d). Unlike the Weibull or Gumbel distributions, there are no values for the location or scale parameters of the Frechet distribution that result in proportional hazard functions.

#### What underlies a unimodal pattern of virulence?

One possibility is for virulence to be correlated with temporal variation in the pathogen’s within-host growth. For example, Bell et al. [26] recorded the growth of *B. bassiana* infections in *An. stephensi* mosquitoes and found the number of conidia detected increased dramatically 3—4 days post-exposure and continued to increase thereafter, but at a slower rate. This increase in conidia replication was correlated with increased fungal division, the growth of hyphae in the host haemocoel, and increased rates of host mortality. These patterns indicate temporal variation in different components of a pathogen’s within-host growth could be correlated with temporal variation in its virulence.

Another means by which unimodal patterns of virulence can arise is when virulence varies among the members of a host population or treatment.

### Variation in virulence

In the examples above virulence varied over time, but the same hazard function described the virulence experienced by all the members of the infected population or treatment. This section considers examples where mortality rates vary among hosts within an infected population or treatment. This variation influences the pattern of mortality observed for the infected population or treatment as a whole and can be qualitatively different from the pattern of virulence experienced by individual hosts.

#### Observed discrete variation

In this example unimodal patterns of virulence were observed for each treatment exposed to infection. However, within each treatment visual cues allowed hosts to be classified into two distinct sub-populations experiencing either virulent or avirulent infections.

Parker et al. [27] followed the survival of uninfected *Acyrthosiphon pisum* pea aphids and those experimentally-infected with the fungal pathogen *Pandora neoaphidis*. The original data are freely available [28]. The experiment involved three dose treatments and six host genotypes. Host genotype is not taken into account in the following analyses, but the effect of dose is; see S11.

Survival was lower in the treatments exposed to infection and tended to decrease in a dose-dependent manner (Fig 5 a-c). Survival in the control treatment suggested that roughly half the mortality observed in infected treatments could be attributed to background mortality. As dose increased, survival curves in the treatments exposed to infection tended to level off indicating mortality rates were slowing over time and suitable for description by the Frechet distribution (Fig 5d-f).

**Figure 5:**
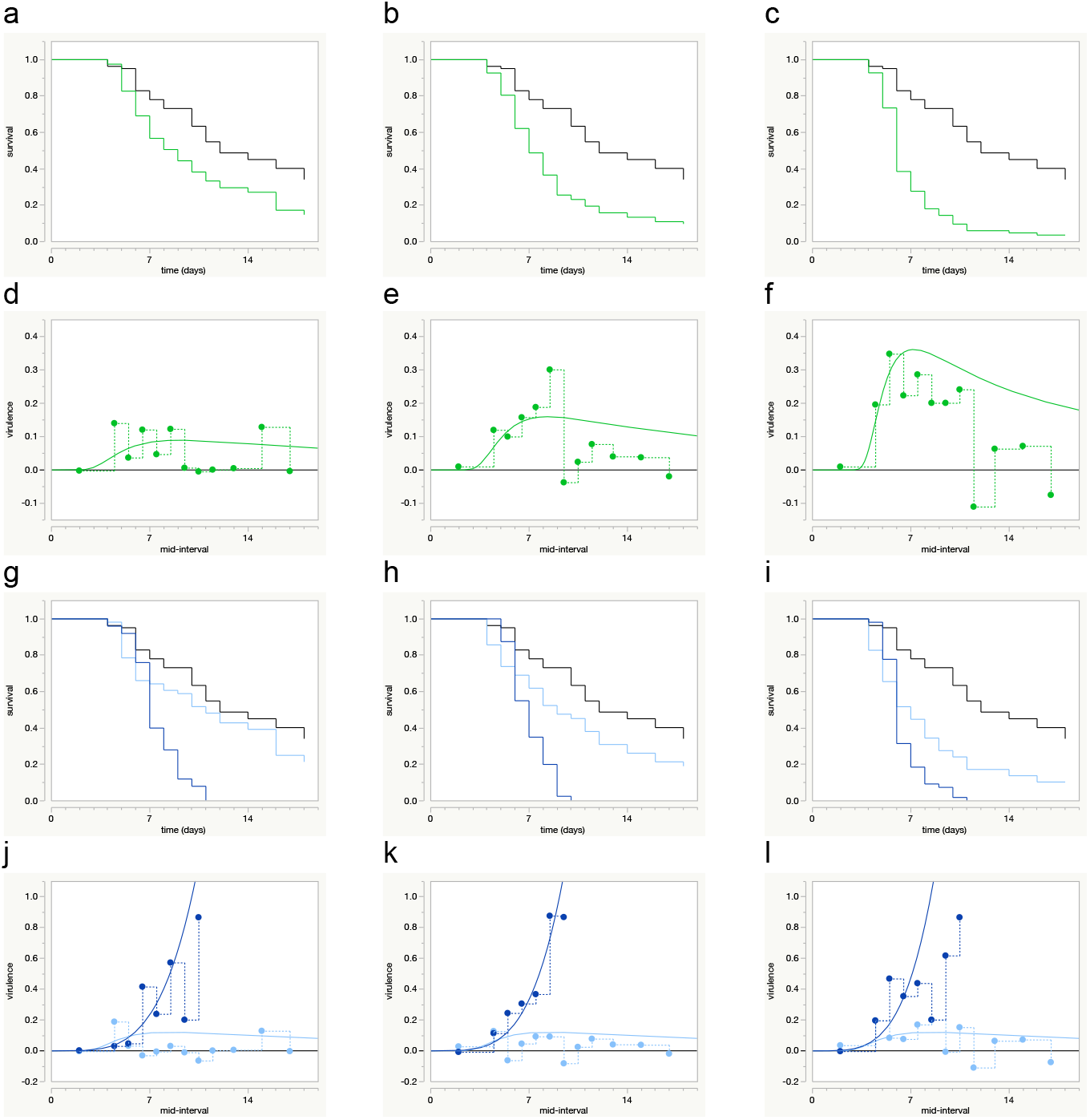
Observed unimodal patterns of virulence and underlying heterogeneity of virulence within populations. Data in the 1st, 2nd and 3rd columns are for aphids exposed to the low, medium and high dose treatments of the fungal pathogen, respectively. (a-c) Observed survival in unexposed control population (black line) and exposed population (green line). (d-f) Observed and estimated unimodal patterns of virulence at the level of the exposed population in each treatment (observed, symbols, dotted stepped line; estimated, smooth curve). (g-i) Observed survival when exposed host population classified according to sporulation status (black, unexposed controls; light blue, non-sporulating; dark blue, sporulating). (j-l) Observed and estimated virulence in sporulating and non-sporulating populations (dark blue symbols/lines, light blue symbols/lines, respectively).

Parker et al. not only recorded when aphids died, but also whether they showed visible signs of fungal sporulation at the time of their death [27]. These individuals accounted for 31%, 49% and 65% of hosts in the low, intermediate and high dose treatments, respectively. The sporulating population all died within 12 days of exposure to infection, whereas those in the non-sporulating population were more likely to survive until the end of the experiment (Fig 5g-i). The log-likelihood model (18) was parameterised such that common location and scale parameters were estimated for the background mortality in each population, but separate location and scale parameters were estimated for mortality due to infection in the sub-populations of sporulating and non-sporulating hosts; see S11.

Observed rates of mortality due to infection increased rapidly over time in the sub-population of sporulating hosts and were described by Weibull hazard functions that accelerated over time (Fig 5j-l). In contrast, the observed and estimated virulence experienced in the sub-population of non-sporulating hosts remained close to zero over time (Fig 5j-l). This suggests these hosts experienced avirulent infections having little or no effect on their rates of mortality.

These results show a monotonically increasing pattern of virulence experienced at the level of individual hosts can appear as a unimodal pattern of virulence at the level of the population as a whole when virulence varies among hosts. These results also show the dose of infection can have qualitatively different effects on a pathogen’s estimated virulence: the analysis of the Lorenz & Koella data [25] suggest virulence was proportional to the dose of infection, whereas the analysis above of the Parker et al. data [28] suggest the main of dose was to influence the proportion of hosts experiencing virulent infections, not the virulence of these virulent infections.

#### Avirulent infections

In the example above, the non-sporulating populations of hosts showed little or no increase in mortality relative to uninfected hosts. This section considers two reasons why hosts in infected treatments may experience avirulent infections

*(i) Exposed-but-uninfected hosts*. Exposure to infection does not guarantee infection. Hosts assumed to be experiencing avirulent infections may have avoided or resisted infection and were never actually infected. If avoiding or resisting infection has no effect on their survival, they should only experience background mortality. In such cases the increased rate of mortality of infected hosts in an infected treatment means they will tend to die earlier than the uninfected members. Consequently the rate of mortality in the treatment as a whole will become increasingly biased towards that for background mortality over time. If the proportions of infected and uninfected members of the population cannot be distinguished and analysed separately, the proportion of exposed-but-uninfected individuals can be estimated from the observed pattern of survival in the population as a whole as,

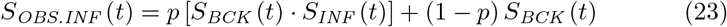

where *p* is a constant to be estimated, 0 ≤ *p* ≤ 1.

Differentiating (23) with respect to time and taking the negative gives the probability density function for mortality observed in the population as a whole at time *t*, *f_OBS.INF_* (*t*), which can be used in turn to estimate the hazard function, *h_OBS.INF_*(*t*); see S12.

This type of model is sometimes referred to as a ‘cure model’ as the observed rate of mortality in the ‘infected’ population will eventually converge with that in a matching uninfected population [29]. However this convergence is due to differential rates of mortality in the infected vs. exposed-but-uninfected populations rather than any process related to hosts being cured or recovering from infection. The following describes a model where ‘avirulent’ infections arise due to hosts recovering from infection.

*(ii) Recovery from infection*. Infected hosts may be able to recover from infection. Here this is assumed to mean they are no longer exposed to the risk of dying due to infection; it does not necessarily mean recovered hosts are uninfected and is not used here in relation to whether hosts are infectious and capable of transmitting disease or not.

This model assumes the pattern of events in an infected population at time *t*, *S_INF.POP_* (*t*), can be described as the product of three independent probability distributions,

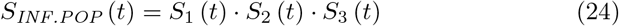

where *S*_1_ (*t*) is the survival function for background mortality at time *t*, *S*_2_ (*t*) is the survival function for mortality due to infection at time *t*, and *S*_3_ (*t*) is the survival function for the probability an infection ‘survives’ until time t, i.e., the host has not recovered at time *t*. Here the index *_INF.POP_*, is used rather than, *_OBS.INF_*, as recovery from infection may not be an observed event.

Differentiating (24) with respect to time and taking the negative gives the probability density function, *f_INF.POP_* (*t*), for events occurring in the population at time *t*,

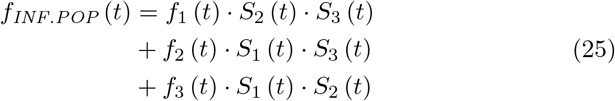

where the sum of the first two expressions gives the probability an infected host is infected and dies at time *t*, from either background mortality or mortality due to infection, and corresponds with data collected on the time of death of infected hosts.

The third expression describes the probability an infected host is alive and recovers from infection at time *t*, and corresponds with data collected on the timing of recovery of infected hosts. Hence the expression can be estimated when the timing of recovery is known (e.g., [30]). This will not be the case if a host’s recovery status is only determined after the host has died or been censored, as the data collected correspond with the time hosts recovered and subsequently survived before dying or being censored. However it is assumed recovered individuals experience the same background mortality as uninfected hosts. This information can be used to estimate the likelihood a recovered individual dying at time t, recovered at an earlier time and subsequently survived until time *t*, when it died or was censored; see S13.

The two types of model are described in greater detail in S12 and S13, respectively, where each model is also applied to the Parker et al. data. These analyses find the exposed-but-uninfected model provided a better description of the data than the recovery model, suggesting the hosts experiencing avirulent infections were more likely to have been uninfected, rather than recovered, hosts.

Code for running the exposed-but-uninfected model in R is given in S12 and that for running the recovery model is described in S13 and provided in a separate file.

#### Unobserved continuous variation

The models above considered virulence when it varied discretely within an infected population or treatment. In this section unobserved variation in mortality rates is assumed to vary continuously among hosts, where all the hosts exposed to infection are infected and there is no recovery from infection.

Frailty models make use of a proportional hazards assumption in which an underlying rate of mortality is multiplied by a constant, e.g. *λ*, where *λ* varies among members of the same population or treatment. Bigger values of this constant are associated with weaker or more ‘frail’ individuals. The distribution of *λ* determines the pattern of survival in the population. In particular, it determines the pattern relative to that expected if mortality rates did not vary among hosts. The difference between the observed and expected patterns of mortality can be estimated as a function of the variance in *λ*, if *λ* is assumed to be distributed as a continuous random variable of a probability distribution constrained to a mean value of one. For example, if unobserved variation in virulence is assumed to follow the gamma distribution with a mean of one, the rate of mortality due to infection in the infected population at time *t*, *h_INF_* (*t*), can be expressed as,

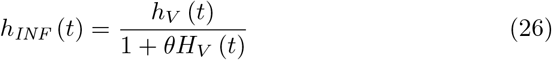

where *h_V_* (*t*) and (*t*) are the hazard and cumulative hazard functions for the underlying pattern of virulence at time *t*, respectively, while *θ* is a constant describing the variance of the distribution of *λ* and a parameter to be estimated.

The average frailty of individuals alive at time *t* in the infected population decreases over time as, [1 + *θH_V_* (*t*)]^−1^. The corresponding function for survival due to infection in the infected population at time *t*, *S_INF_* (*t*), is,

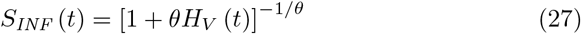

These expressions were used to analyse the data from the Lorenz & Koella study [24], when data from all six dose treatments were pooled to form a single infected population. This population was thus known to be heterogeneous for virulence. Furthermore as virulence was estimated as being proportional to dose, the proportional relationship assumed between the underlying pattern of virulence and that experienced by individual hosts should have been met.

Although the pattern of virulence within individual dose treatments generally increased monotonically over time (c.f. Fig 3b), the pattern for the infected population as a whole was unimodal (Fig 6a). This unimodal pattern was captured by a likelihood model where the hazard function of the Frechet distribution described the pathogen’s virulence (Fig 6a). This model (incorrectly) assumed virulence was homogenous among infected hosts; see S14.

**Figure 6:**
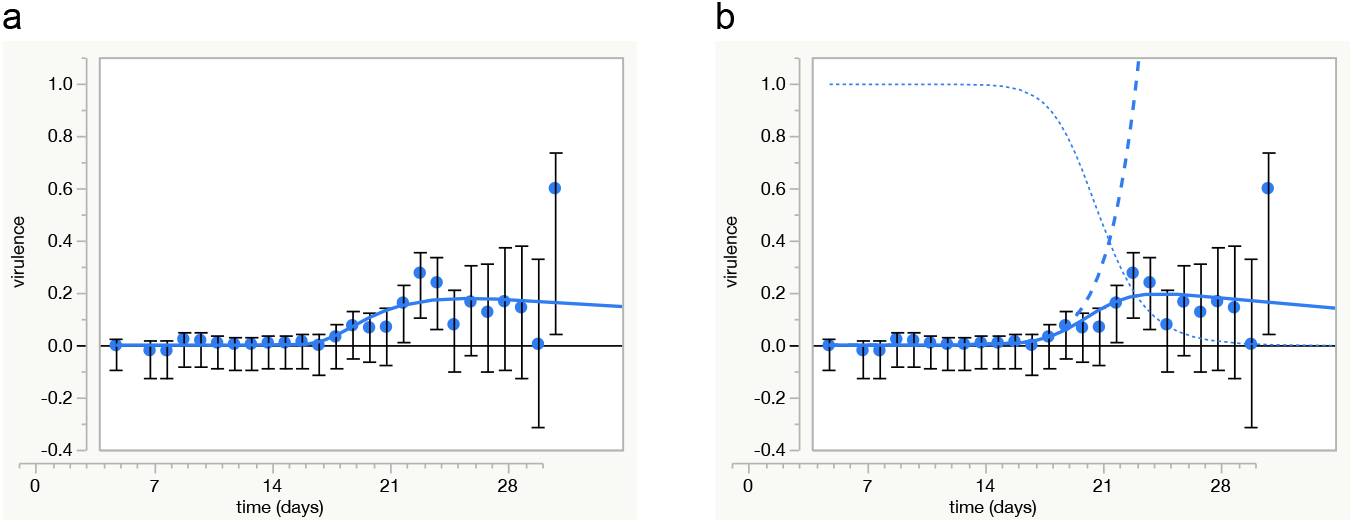
Unobserved variation in virulence with a continuous distribution. (a) Daily estimates of virulence from life table data for the pooled population of infected female mosquitoes (symbols ±95%*c.i*.). The blue line is a Frechet hazard function estimating virulence, (b) frailty model estimates for underlying pattern of virulence experienced by individual hosts (dashed line) and average virulence experienced at level of the infected population (solid line), along with the average frailty of infected individuals alive at time *t* (dotted line) based on Weibull hazard functions and gamma distributed frailty.

The unimodal pattern of virulence experienced at the level of the population was also captured by a frailty model (Fig 6b). This model (correctly) assumed virulence varied among infected hosts. The Weibull hazard function describing the underlying pattern of virulence experienced by individual hosts increased over time (Fig 6b). However this increase was counteracted at the level of the population by the progressive decrease in the average frailty of the infected hosts remaining alive (Fig 6b).

The frailty model above only allowed for unobserved variation in mortality due to infection. It could have been considered as acting on background mortality or on both sources of mortality in a more or less correlated fashion; see Zahl [31] and S15 for details.

Frailty models can also be adapted to allow for random variation at higher levels of organisation than considered above, e.g., among replicate populations of the same treatment or among groups of hosts sharing the same genotype. These multivariate frailty models are more complicated to formulate and are beyond the scope of this study. For more information on multivariate models and frailty models in general see; Hougaard [32, 33], Aalen [34], Gutierrez [35].

### Virulence and transmission

Epidemiological models generally assume a pathogen’s virulence arises as an unavoidable consequence of its within-host growth and production of transmission stages [1–3]. Empirical studies estimating a pathogen’s virulence can investigate this relationship when they also estimate the pathogen’s transmission success.

Empirical estimates of transmission success often rely on measures of the pathogen’s potential, rather than its actual, transmission success. For example, Sy et al. [36, 37] estimated the potential transmission success of *V. culicis* from the number of the pathogens’ spores adult *Aedes aegypti* female mosquitoes harboured at the time of their death. These spores are not shed while the host is alive and thus represent the pathogen’s cumulative investment into its potential transmission success over its host’s lifetime. This cumulative sum was positively correlated with estimates for the host’s cumulative exposure to the pathogen’s virulence, *H_INF_* (*t*) (Fig 7a). This could suggest a causal relationship between the production and accumulation of the pathogens’ spores and its virulence, but this correlation is potentially confounded as the host’s exposure to background mortality, *H_BCK_* (*t*), also increased over time in a similar manner. However the number of spores hosts harboured was positively correlated with the ratio, *H_INF_* (*t*)/*H_OBS.INF_* (*t*), at the time hosts died (Fig 7b). This ratio varies between 0 and 1 and reflects the estimated contribution of the pathogen’s virulence towards the host’s overall exposure to the risk of dying at the time they died. Thus when hosts died in the study by Sy et al. [36], this ratio was generally greater than 0.5 and positively correlated with the number of spores they harboured, suggesting the cumulative effects of infection and spore production were positively correlated with host mortality.

**Figure 7:**
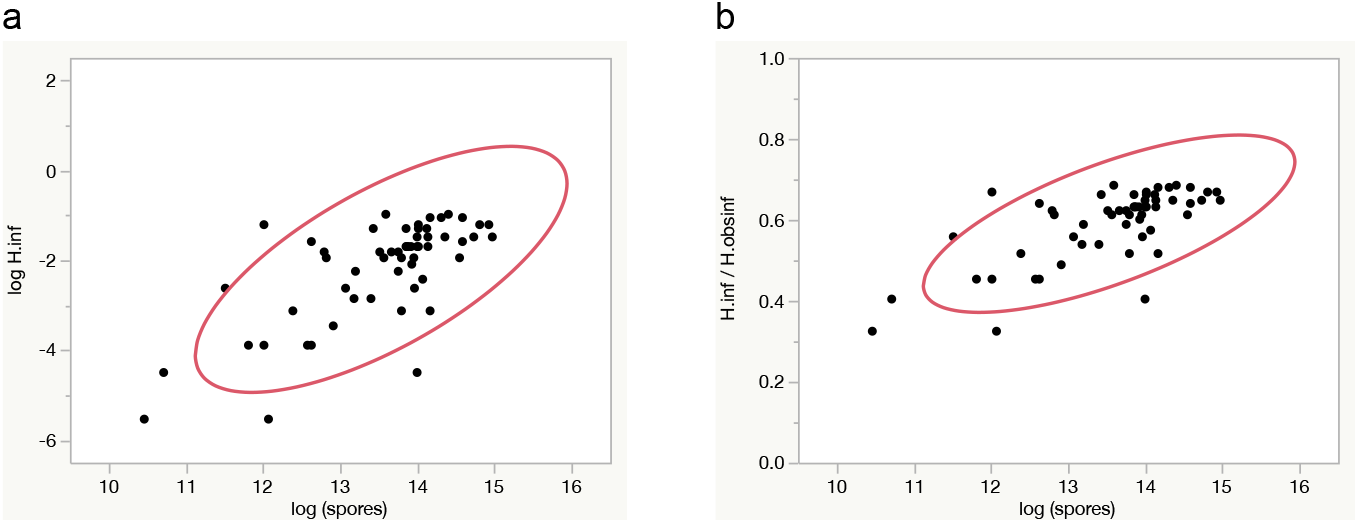
Correlations between cumulative spore production and cumulative virulence. The log (number of *V. culicis* spores) recovered from adult mosquitoes at the time of their death was positively correlated with estimates for, (a) the host’s cumulative exposure to the pathogen’s virulence, *H_INF_* (*t*), at the time of their death and (b) the contribution of *H_INF_* (*t*) to the host’s cumulative exposure to the risk of dying, *H_INF_* (*t*)/*H_OBS.INF_* (*t*), at the time of their death.

#### Comparing estimates

It is often convenient to compare estimates of virulence and transmission success at a particular point in time. For example, pathogen fitness as estimated by *R*_0_ is often calculated based on the estimated time infected hosts are infectious [6, 7].

When there is no recovery from infection, the average longevity of infected hosts can be estimated by integrating the survival function *S_OBS.INF_* (*t*). However if host rates of mortality are constant, the average longevity of infected hosts conveniently equals, 1/(*μ* + *ν*). It also corresponds with the time when *H_OBS.INF_* (*t*) = 1, which is the time when infected hosts are expected to have died due to their cumulative exposure to the risk of dying from background mortality and mortality due to infection (9).

When host mortality rates vary over time, and there is no recovery from infection, the average longevity of infected hosts and the time when *H_OBS.INF_* (*t*) = 1 are not the same. However it may be convenient to use the time when *H_OBS.INF_* (*t*) = 1 for comparative purposes as it can be directly read from plots of cumulative survival in infected treatments: when *H_OBS.INF_* (*t*) = 1, *S_OBS.INF_* (*t*) = exp(−1) = °.368 (5).

When *S_OBS.INF_* (*t*) = 0.367, the cumulative survival curve in a matching control treatment can be used to estimate the host’s cumulative exposure to background mortality,

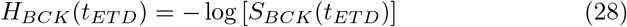

and thus its cumulative exposure to the pathogen’s virulence,

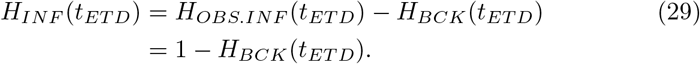

where *t_ETD_* is the ‘expected time of death’ for infected hosts when *H_OBS.INF_* (*t*) = 1. These estimates do not require any assumptions as to the distribution of background mortality or mortality due to infection.

For example, the cumulative survival curves for infected hosts in the high and low dose treatments of the study by Lorenz & Koella [24] intersected the line, *S_OBS.INF_* (*t*) = 0.368, on days 17.5 and 21.5, respectfully (Fig 8). At these times, survival in the control treatment was 0.784 and 0.619, respectively. The negative log-transformation of these values estimated the host’s cumulative exposure to the risk of dying from background mortality, *H_BCK_*(*t_ETD_*), as 0.243 and 0.480, respectively, giving estimates of the host’s cumulative exposure to the pathogen’s virulence at its expected time of death, (*t_ETD_*), as, 0.757 and 0.520, respectively.

**Figure 8:**
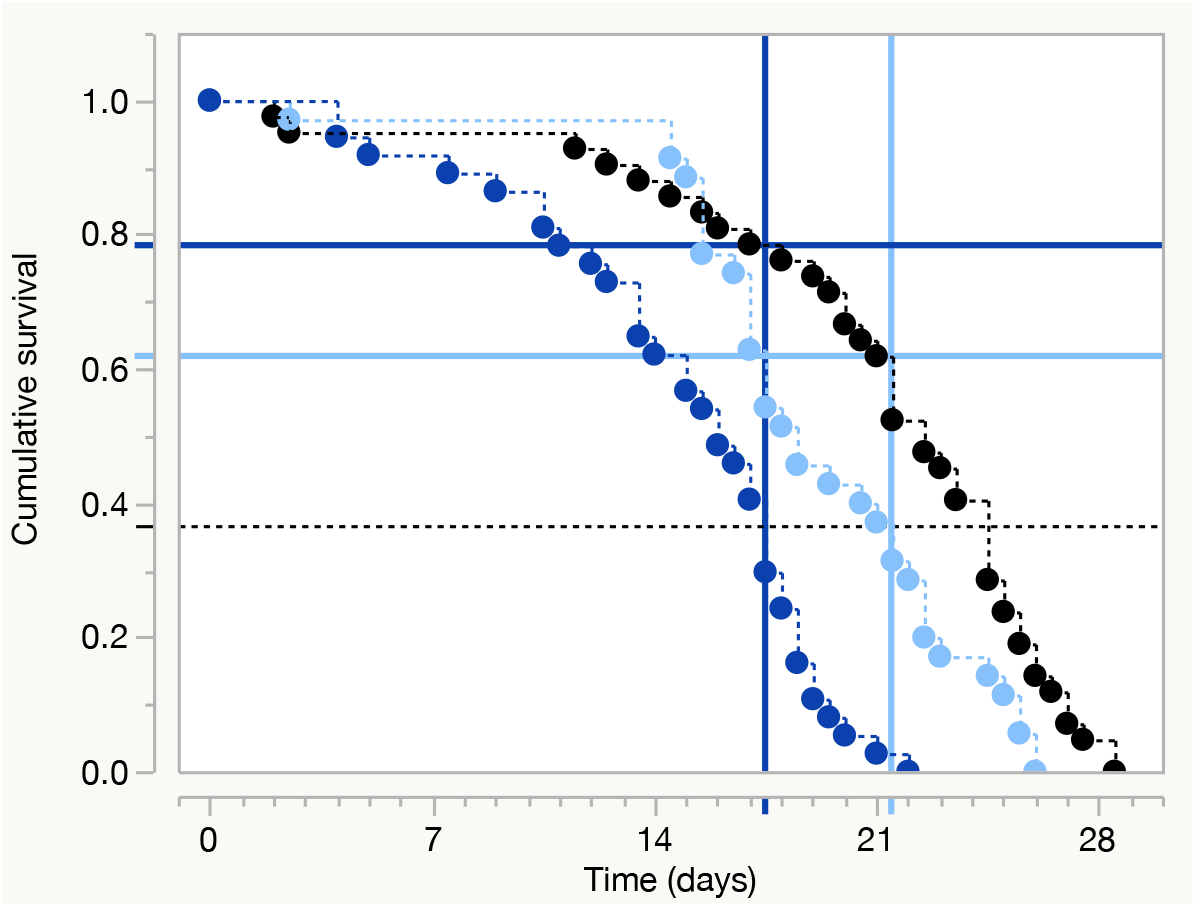
Direct estimates of cumulative virulence from survival curve data. Cumulative survival in the control (black), low dose (light blue) and high dose (dark blue) treatments of the study by Lorenz & Koella. The horizontal dotted line is when *H_OBS.INF_* (*t*) = 1 and *S_OBS.INF_* (*t*) = 0.368. The intersection of this line with the cumulative survival curves in the infected treatments give the ‘expected times of death’ for infected hosts in the infected treatments (vertical blue lines). The intersection of these vertical lines with the cumulative survival curve for the control treatment estimates the contribution of background mortality to the observed survival of infected hosts (horizontal blue lines). These values allow estimation of the contribution of background mortality and the pathogen’s virulence to the survival of infected hosts at their ‘expected time of death’. Re-drawn from Fig. 3a with data pooled across larval food treatments.

Hence infected hosts were not only expected to die earlier in the highest vs. lowest dose treatment (17.5 vs. 21.5 days), but their death was estimated as more likely to be due to infection (0.757 vs. 0.520). When hosts in the lowest dose treatment were expected to die, the likelihood of them dying from infection was approximately equal to that of them dying from background mortality (0.520 vs. 0.480), whereas it was roughly three times greater in the highest dose treatment (0.757 vs 0.243).

### Wider applications of relative survival

The analysis of relative survival can be applied when survival in a target population is less than would be expected due to a specific factor. These situations extend beyond those involving patients afflicted with a particular disease or illness or hosts experimentally-infected with a particular pathogen. The analysis of relative survival could, for example, be usefully applied to estimating the adverse effects of xenobiotics (antibiotics, insecticides, fungicides, etc…) on survival in treated vs. untreated population, such as, populations of bees exposed or unexposed to neonicotinoid pesticides, or bees experimentally exposed to different combinations of stressors, such as, pesticides and pathogens.

Another application of the method could be to compare survival in model organisms, such as, *Drosophila melanogaster* or *Caenorhabditis elegans*, where their survival is reduced following the knockdown of a particular gene, gene-product, or component of the immune system, etc.…Adopting the analysis of relative survival in these studies could provide much more detailed estimates for the effects of the experimental manipulation than the log-rank tests habitually used to test for evidence of effect.

## Summary

Reciprocal feedback between data and theory is one of the motors by which science advances. This progress is potentially hindered when empirical and theoretical studies differ in how the subject of interest is defined. This can mean like is not being compared with like and the assumed relationship between two variables is potentially confounded by a third variable [4].

Most empirical studies generating estimates of a pathogen’s virulence do not estimate virulence as it is defined in the theoretical literature [1–3], where it is generally defined as the increase in the per capita rate of mortality of infected hosts due to infection [6, 7]. The aim of this study has been to illustrate how this rate can be estimated from the data of empirical studies by applying a statistical method widely used in the medical literature to analyse relative survival in populations of patients afflicted by a particular disease or illness [8–11]. This does not validate the theoretical literature’s use of this definition for virulence, but it allows theoretical and empirical studies to compare virulence based on the same metric.

Worked examples illustrate cases where rates of host mortality due to infection accelerate over time and where this acceleration is proportional to the dose of infection to which hosts were exposed. Other examples illustrate the observed pattern of virulence in an infected population or treatment can increase and then decrease over time. This unimodal pattern could reflect the pattern of the pathogen’s within-host growth, but can also indicate virulence varied among members of the host population. Such variation may be directly observed or assumed to be present and analysed as discretely or continuously distributed. These estimates open the way for the relationship between virulence and other traits to be explored, such as, transmission success, and how they contribute to pathogen fitness.

## Supporting information

Blanford et al data

R recovery model and data

## Acknowledgements

I thank Simon Blanford for generously providing data upon request and Matthew Thomas for permission to release the data. I also thank Yannis Michalakis for according me the time to complete this study and comments on earlier versions of the manuscript. Eric Elguero provided useful input during discussions on recovery from infection.

This work was funded by basic research funds from the French research agencies of the Centre National de la Recherche Scientifique (CNRS) and the Institut de Recherche pour le Développement (IRD).

## References

1. Alizon, S., Hurford, A., Mideo, N. & van Baalen, M. Virulence evolution and the trade-off hypothesis: history, current state of affairs and the future. Journal of Evolutionary Biology 22, 245–259 (2009).

2. Schmid-Hempel, P. Evolutionary Parasitology: The Integrated Study of Infections, Immunology, Ecology, and Genetics (OUP Oxford, 2011).

3. Cressler, C. E., McLeod, D. V., Rozins, C., van den Hoogen, J. & Day, T. The adaptive evolution of virulence: a review of theoretical predictions and empirical tests. Parasitology 143, 915–930 (2016).

4. Day, T. On the evolution of virulence and the relationship between various measures of mortality. Proceedings of the Royal Society B-Biological Sciences 269, 1317–1323 (2002).

5. Read, A. F. The evolution of virulence. Trends in Microbiology 2, 73–76 (1994).

6. Anderson, R. M. & May, R. M. Population biology of infectious-diseases. 1. Nature 280, 361–367 (1979).

7. May, R. M. & Anderson, R. M. Population biology of infectious-diseases. 2. Nature 280, 455–461 (1979).

8. Ederer, F., Axtell, L. M. & Cutler, S. J. The relative survival rate: a statistical methodology. Natl Cancer Inst Monogr 6, 101–121 (1961).

9. Monson, R. R. Analysis of relative survival and proportional mortality. Computers and Biomedical Research 7, 325–332 (1974).

10. Esteve, J., Benhamou, E., Croasdale, M. & Raymond, L. Relative survival and the estimation of net survival - elements for further discussion. Statistics in Medicine 9, 529–538 (1990).

11. Dickman, P. W., Sloggett, A., Hills, M. & Hakulinen, T. Regression models for relative survival. Statistics in Medicine 23, 51–64 (2004).

12. Chow, W. H., Devesa, S. S., Warren, J. L. & Fraumeni, J. F. Rising incidence of renal cell cancer in the United States. Jama-Journal of the American Medical Association 281, 1628–1631 (1999).

13. Hundahl, S. A., Fleming, I. D., Fremgen, A. M. & Menck, H. R. A National Cancer Data Base Report on 53,856 Cases of Thyroid Carcinoma Treated in the US, 1985-1995. Cancer 83, 2638–2648 (1998).

14. Brown, L. & Li, X. F. Confidence intervals for two sample binomial distribution. Journal of Statistical Planning and Inference 130, 359–375 (2005).

15. Agresti, A. & Caffo, B. Simple and effective confidence intervals for proportions and differences of proportions result from adding two successes and two failures. American Statistician 54, 280–288 (2000).

16. Austin, P. C. A Tutorial on Multilevel Survival Analysis: Methods, Models and Applications. International Statistical Review 85, 185–203 (2017).

17. Hintze, J.L. NCSS User’s Guide V. Chapter 550, Distribution (Weibull) Fitting NCSS (Kaysville, Utah, 2007). https://www.ncss.com/.

18. Hurvich, C. M. & Tsai, C. L. Regression and time-series model selection in small samples. Biometrika 76, 297–307 (1989).

19. R Core Team. R: A Language and Environment for Statistical Computing R Foundation for Statistical Computing (Vienna, Austria, 2017). https://www.R-project.org/.

20. Bolker, B. and R Development Core Team. bbmle: Tools for General Maximum Likelihood Estimation R package version 1.0.19 (2017). https://CRAN.R-project.org/package=bbmle.

21. SAS Institute Inc. JMP@ 13 (Cary, NC, 2016).

22. Blanford, S., Jenkins, N. E., Read, A. F. & Thomas, M. B. Evaluating the lethal and pre-lethal effects of a range of fungi against adult Anopheles stephensi mosquitoes. Malaria Journal 11, 10 (2012).

23. Cox, D. R. Regression models and life-tables. Journal of the Royal Statistical Society Series B-Statistical Methodology 34, 187–220 (1972).

24. Lorenz, L. M. & Koella, J. C. The microsporidian parasite Vavraia culicis as a potential late life-acting control agent of malaria. Evolutionary Applications 4, 783–790 (2011).

25. Lorenz, L. M. & Koella, J. C. Data from: The microsporidian parasite Vavraia culicis as a potential late life-acting control agent of malaria. Dryad Digital Repository. doi:10.5061/dryad.2s231 (2011).

26. Bell, A. S., Blanford, S., Jenkins, N., Thomas, M. B. & Read, A. F. Real-time quantitative PCR for analysis of candidate fungal biopesticides against malaria: Technique validation and first applications. Journal of Invertebrate Pathology 100, 160–168 (2009).

27. Parker, B. J., Garcia, J. R. & Gerardo, N. M. Genetic variation in resistance and fecundity tolerance in a natural host-pathogen interaction. Evolution 68, 2421–2429 (2014).

28. Parker, B. J., Garcia, J. R. & Gerardo, N. M. Data from: Genetic variation in resistance and fecundity tolerance in a natural host-pathogen interaction. Dryad Digital Repository. doi:10.5061/dryad.24gq7 (2014).

29. Lambert, P. C. Modeling of the cure fraction in survival studies. Stata Journal 7, 351–375 (2007).

30. Gervasi, S., Burgan, S. C., Hofmeister, E., Unnasch, T. R. & Martin, L. B. Stress hormones predict a host superspreader phenotype in the West Nile virus system. Proceedings of the Royal Society of London B: Biological Sciences 284. doi:10.1098/rspb.2017.1090 (2017).

31. Zahl, P. H. Frailty modelling for the excess hazard. Statistics in Medicine 16, 1573–1585 (1997).

32. Hougaard, P. Life table methods for heterogeneous populations - distributions describing the heterogeneity. Biometrika 71, 75–83 (1984).

33. Hougaard, P. Frailty models for survival data. Lifetime Data Analysis 1, 255–273 (1995).

34. Aalen, O. O. Heterogeneity in survival analysis. Statistics in Medicine 7, 1121–1137 (1988).

35. Gutierrez, R. G. Parametric frailty and shared frailty survival models. Stata Journal 2, 22–44 (2002).

36. Sy, V. E., Agnew, P., Sidobre, C. & Michalakis, Y. Reduced survival and reproductive success generates selection pressure for the dengue mosquito Aedes aegypti to evolve resistance against infection by the microsporidian parasite Vavraia culicis. Evolutionary Applications 7, 468–479 (2014).

37. Sy, V. E., Agnew, P., Sidobre, C. & Michalakis, Y. Data from: Reduced survival and reproductive success generates selection pressure for the dengue mosquito Aedes aegypti to evolve resistance against infection by the microsporidian parasite Vavraia culicis. Dryad Digital Repository. doi:10.5061/dryad.1rs10 (2014).

