## Supplementary files for "Estimating virulence from relative survival"

S01. Estimating virulence & confidence intervals at individual points in time
S02. Delta method for calculating confidence intervals of a hazard function
S03. Maximum likelihood estimation of virulence using R
S04. Maximum likelihood estimation of virulence using JMP
S05. Initial parameter values
S06. Analysis of Blanford et al. data (i)
S07. Estimating average longevity in R
S08. Analysis of Lorenz & Koella data (i)
S09. Accelerated failure time (AFT) model
S10. Analysis of Blanford et al. data (ii)
S11. Analysis of Parker et al. data
S12. Exposed-but-uninfected hosts model
S13. Recovery from infection model
S14. Lorenz & Koella pooled data
S15. Shared and correlated frailty models

### S01 Estimating virulence & confidence intervals at individual points in time

#### Regular sampling intervals

When host mortality is sampled at regular intervals, the rate of mortality in interval $i$, $h(i)$ can be estimated as,

$$h(i) = d(i)/r(i)$$

where $d(i)$ is the number of individuals dying in the interval $i$ and $r(i)$ is the number of individuals alive or at risk of dying at the beginning of interval $i$.

In the context of relative survival, the observed rate of mortality in an infected treatment, $h_{OBS.INF}(t)$, is the sum of the rate of background mortality, $h_{BCK}(t)$, plus the rate of mortality due to infection, $h_{INF}(t)$. Consequently, the difference in the two observed rates of mortality

$$h_{INF}(t) = h_{OBS.INF}(t) - h_{BCK}(t)$$

provides an estimate of the pathogen's virulence at time $t$. This is the difference between two binomial proportions. Following Agresti & Caffo (1), if $p_1$ is the proportion of infected individuals observed dying in the interval and $p_2$ the proportion of uninfected individuals dying in the same interval, the 95% confidence intervals for the adjusted Wald interval of $p_1 - p_2$, can be calculated as,

$$(\tilde{p}_1 - \tilde{p}_2) \pm 1.96\sqrt{\frac{\tilde{p}_1(1-\tilde{p}_1)}{\tilde{n}_1} + \frac{\tilde{p}_2(1-\tilde{p}_2)}{\tilde{n}_2}}$$

where $\tilde{p}_i = (X_i + 2)/(n_i + 4)$ and $\tilde{n}_i = (n_i + 4)$, $X_i$ is the number of individuals dying in the interval and $n_i$ the number of individuals at risk of dying at the beginning of the interval, for $i = 1, 2$ respectively. The value of $+2$ is an approximation for the $Z$ score of 1.96 for the 97.5 percentile used when calculating 95% confidence intervals, $+4$ an approximation for $1.96^2$.

#### Irregular sampling intervals

If intervals among sampling times are irregular, the calculations above need correcting to standardise the rate over which virulence is estimated. One way to do this is to divide the number of individuals dying during the interval $(d)$ by the duration or breadth of the interval $(b)$, and dividing the result by the expected number of individuals alive at the midpoint $(t_{mt})$ of the interval,

$$h(t_{mt}) = \frac{d/b}{r - d/2}$$

where $r$ is the number of individuals alive, or at risk of dying, at the beginning of the interval and $b$ is the width or duration of the shortest interval. NB the width of the shortest interval requires $b \geq 1$ to avoid estimates of $h(t_{mt}) > 1$. For example if the shortest interval between samples was 0.5 days, it would be better to calculate rates of mortality in units of 12 hours, rather than 0.5 days.

(1) doi:10.2307/2685779

#### S02 Delta method for calculating confidence intervals of a hazard function

This approach to estimating confidence intervals for a hazard function is based on the delta method and is taken from *Chapter 550 Distribution (Weibull) Fitting for the NCSS Statistical Software, pg. 268* (1), which in turn cites, *Nelson WB. 1990. Accelerated Testing. John Wiley, New York (pg. 294).*

The confidence intervals for an estimated hazard function at time $t$, $\hat{h}(t)$, are

$$\hat{h}\left(t\right)\exp\left[\pm\frac{z_{1-\alpha/2}s\left[\hat{h}\left(t\right)\right]}{\hat{h}\left(t\right)}\right]$$

where

$$s^2\left[\hat{h}\left(t\right)\right]=\left(\frac{\partial\hat{h}}{\partial a}\right)^2\operatorname{var}\left(a\right)+\left(\frac{\partial\hat{h}}{\partial b}\right)^2\operatorname{var}\left(b\right)+2\left(\frac{\partial\hat{h}}{\partial a}\right)\left(\frac{\partial\hat{h}}{\partial b}\right)\operatorname{cov}\left(a,b\right)$$

$a$ and $b$ are the location and scale parameters, var($a$) and var($b$) are the variances of estimates of $a$ and $b$, and cov($a$,$b$) their covariance, respectfully.

For example, the partial derivatives for the Weibull hazard function,

$$h\left(t\right)=\left(\frac{1}{bt}\right)\exp\left(z\right)$$

where $z=\left(\log t-a\right)/b$ are,

$$\frac{\partial h}{\partial a}=-\frac{1}{b^2t}\exp\left(t\right)$$

and

$$\frac{\partial h}{\partial b}=-\frac{1}{b^2t}\left[z\exp\left(z\right)-\exp\left(z\right)\right]$$

(1) http://www.ncss.com/wp-content/uploads/2012/09/NCSSUG5.pdf

### S03 Maximum likelihood estimation of virulence using *R*

This supplement provides code specifying the log-likelihood expression allowing the location and scale parameters for background mortality and mortality due to infection to be estimated by maximum likelihood using the package *bbmle* (1) for *R* (2).

#### Individual data

The model is specified for a table with survival data in the format;

| Individual | $t$ | $d$ | $g$ |
|---|---|---|---|
| 1 | 10 | 0 | 1 |
| 2 | 12 | 1 | 0 |
| ⋮ | | | |
| n | 11 | 1 | 1 |

where there is a row of data for each of the $n$ individuals to be analysed;
$t$ is the time when the individual died or was right-censored,
$d$ is a death indicator variable taking a value of '1' for individuals that died and '0' for those censored,
$g$ is an infection indicator variable taking a value of '1' for individuals in the infected treatment and '0' for those in the uninfected control treatment.

The log-likelihood expression to be evaluated is,

$$\log L = \sum_{i=1}^{n} \{\ d \log\left[h_{BCK}(t_i) + g h_{INF}(t_i)\right] + \log\left[S_{BCK}(t_i)\right] + g \log\left[S_{INF}(t_i)\right]\ \}$$

where $h_{BCK}(t)$, $h_{INF}(t)$, $S_{BCK}(t)$ and $S_{INF}(t)$ are the hazard and survival functions for background mortality and mortality due to infection for individual $i$ at time $t$, respectively; $d$ and $g$ are defined as above.

#### Model

```
LL01 <- function(au,bu,ai,bi,datatable){

        zu <- (log(t) - au)/bu
        zi <- (log(t) - ai)/bi

        hu <- (1/(bu*t))*exp(zu)
        hi <- (1/(bi*t))*exp(zi)

        Su <- exp(-exp(zu))
        Si <- exp(-exp(zi))

        logl <- -sum(d*log(hu+g*hi)+log(Su)+g*log(Si))
}
```

```
m01 <- mle2(
  LL01,
  start=list(au=2,bu=0.5,ai=2,bu=0.5,
  data=datatable)

summary(m01)
```

The text defines the function *LL01* which is the log-likelihood expression for analysing relative survival. Here the model is specified using Weibull distribution functions to describe the background mortality and mortality due to infection.

The 1st line defines the parameters to be estimated; here the location and scale parameters *au* and *bu* for the background mortality and *ai*, *bi* for mortality due to infection, respectively, as well as identifying the *datatable* containing the data.

The 2nd and 3rd lines define the $z$ terms corresponding with the Weibull distribution for background mortality ($zu$), and mortality due to infection, ($zi$).

The 4th and 5th lines define Weibull hazard functions for background mortality and mortality due to infection, respectively, making use of the $z$ terms defined above.

The 6th and 7th lines define Weibull survival functions for background mortality and mortality due to infection, respectively, making use of the $z$ terms defined above.

The 8th line is the negative log-likelihood expression using the terms defined above and the indicator variables $d$ and $g$ to be found in the *datatable.*

The maximum likelihood estimation is performed by the *mle2* function of the package *bbmle* (2) and requires the initial values of *au*, *bu*, *ai* and *bi* to be specified.

The results of the analysis are held by *m01.*

#### Grouped data

Instead of there being a single row for each individual, data may be grouped into the frequency of events occurring at a particular time, e.g., in Row 2 of the table below, 15 uninfected individuals ($g = 0$) died ($d = 1$) when $t = 12$.

| Individual | t | d | g | fq |
|---|---|---|---|---|
| 1 | 10 | 0 | 1 | 5 |
| 2 | 12 | 1 | 0 | 15 |
| ⋮ | | | | |
| N | 11 | 1 | 1 | 7 |

In this case the terms in the log-likelihood expression need multiplying by '*fq*';

```
logl <- -sum(fq*(d*log(hu + g*hi) + log(Su) + g*log(Si)))
```

#### Model refinements

##### (i) Making the location parameter for mortality due to infection a function of three dose treatments (*d1, d2, d3*)

This can be achieved by creating a dummy variable for each treatment which takes a value of '1' when the treatment corresponds with that the individual experienced and '0' otherwise, e.g.,

```
id.d1 <- ifelse(datatable$dose=="d1",1,0)
id.d2 <- ifelse(datatable$dose=="d2",1,0)
id.d3 <- ifelse(datatable$dose=="d3",1,0)
```

To quantify the effect of these three dose treatments requires the estimation of two additional parameters, e.g., *aid1* and *aid2*. These are added to the function defining the log-likelihood expression in the first line.

```
LL02 <- function(au,bu,ai,bi,aid1,aid2,datatable){

aid <- ai + aid1*id.d1 + aid2*id.d2
        -(aid1+aid2)*id.d3

         zu <- (log(t) - au)/bu
         zi <- (log(t) - aid)/bi
         hu <- (1/(bu*t))*exp(zu)
         hi <- (1/(bi*t))*exp(zi)
         Su <- exp(-exp(zu))
         Si <- exp(-exp(zi))

         logl <- -sum(d*log(hu + g*hi) + log(Su) + g*log(Si))
 }

m02 <- mle2(LL02,start=list(au=2,bu=0.5,ai=2,bi=0.5,
        aid1=0,aid2=0), data=datatable)
```

```
summary(m02)
```

Here *aid* is created for the effect of dose on the location parameter of mortality due to infection. It estimates the underlying value of the location parameter *ai* plus the deviation due to dose treatment *d1*, *+aid1*id.d1*, and that due to dose treatment *d2*, *+aid2*id.d2*. The deviation due to dose treatment *d3* is the negative of the sum of the other parameters multiplied by the dummy variable; *-(aid1 + aid2)*ai.d3*

The terms *ai* can now be replaced by *aid*. In this example, this only involves the *zi* term. Had the effect of dose on the scale parameter been defined in a similar manner, the terms *bi* in the terms *zi* and *hi* would have needed replacing with *bid*.

The difference between the results of these two models can then be compared in terms of their AIC values

```
AIC(m01,m02)
```

or by likelihood ratio tests

```
anova(m01,m02)
```

The variance and covariance of the estimated parameters needed for the calculation of confidence intervals via the delta method

```
vcov(m01)
```

**(ii) To specify the Gumbel distribution**

```
zx <- (t - ax)/bx
hx <- (1/bx)*exp(zx)
Sx <- exp(-exp(zx))
```

where $x$ is replaced by either $u$ or $i$ when describing the background mortality or mortality due to infection, respectively.

**(iii) To specify the Fréchet distribution**

```
zx <- (log(t) - ax)/bx
hx <- (1/(bx*t))*exp(-zx-exp(-zx))/(1-exp(-exp(-zx))
Sx <- 1-exp(-exp(-zx))
```

where $x$ is replaced by either $u$ or $i$ when describing the background mortality or mortality due to infection, respectively.

Alternatively,

```
zx <- (log(t) - ax)/bx
fx <- (1/(bx*t))*exp(-zx-exp(-zx))
Sx <- 1-exp(-exp(-zx))
hx <- fx/Sx
```

(1) https://CRAN.R-project.org/package=bbmle
(2) https://www.R-project.org/

### S04 Maximum likelihood estimation of virulence using JMP

The figure below is from the formula builder of JMP (1) showing how a log-likelihood expression for analysing relative survival can be specified.

$$\mathrm{zu} = \frac{\left(\mathrm{Log}\left(\mathrm{t}\right) - \mathrm{a0}\right)}{\mathrm{b0}};$$

$$\mathrm{bu} = \mathrm{b0};$$

$$\mathrm{zi} = \frac{\left(\mathrm{Log}\left(\mathrm{t}\right) - \mathrm{a1}\right)}{\mathrm{b1}};$$

$$\mathrm{bi} = \mathrm{b1};$$

$$-\left(\begin{array}{l} \mathrm{d} \bullet \mathrm{Log}\left(\left(\frac{1}{\left(\mathrm{bu} \bullet \mathrm{t}\right)}\right) \bullet \mathrm{Exp}\left(\mathrm{zu}\right) + \mathrm{inf} \bullet \left(\frac{\left(\left(\frac{1}{\left(\mathrm{bi} \bullet \mathrm{t}\right)}\right) \bullet \mathrm{Exp}\left(-\mathrm{zi} - \mathrm{Exp}\left(-\mathrm{zi}\right)\right)\right)}{\left(1 - \mathrm{Exp}\left(-\mathrm{Exp}\left(-\mathrm{zi}\right)\right)\right)}\right)\right) \\ + -\mathrm{Exp}\left(\mathrm{zu}\right) \\ + \mathrm{inf} \bullet \mathrm{Log}\left(1 - \mathrm{Exp}\left(-\mathrm{Exp}\left(-\mathrm{zi}\right)\right)\right) \end{array}\right);$$

Figure S4.1: Screen shot of log-likelihood expression for analysis of relative survival with non-linear platform of JMP

The first four expressions from the top are local variables defined in terms of time, $\log t$, and the four location and scale parameters to be estimated; *a0*, *b0*, *a1*, *b1*.

The last term is the negative log-likelihood expression where the terms describing the background mortality are specified according to the Weibull distribution while those for mortality due to infection are specified for the Fréchet distribution. *d* refers to a column in the data table indicating whether individuals died during the experiment or were right-censored, with values 1 and 0, respectively. inf also refers to a column in the data table and has values of 1 or 0 for infected and uninfected individuals, respectively.

To run the model, follow the pathway; *Analyze/Modeling/Nonlinear*
this will open the nonlinear fit window. Choose the column containing the log-likelihood expression and click on the ‘Loss’ button to select the model. There is also the option to identify a column in the data table containing the frequency of events (‘Freq’).
Click the ‘OK’ button to open the Nonlinear Fit window.
Click to tick the ‘Loss is Neg LogLikelihood’ box if not already ticked.
Specify the initial values for the parameters to be estimated.
Click ‘Go’.
The model will then go through an iterative process until it converges on parameter values that give the best fit between the model and the data.

(1) SAS Institute Inc. (2015) Using JMP 12. Cary, NC.

#### S05 Initial parameter values

Maximum likelihood estimation techniques used to solve likelihood problems require initial values for the parameters to be estimated. These are progressively adjusted in an iterative process until the fit between the likelihood model and the observed data does not improve beyond a threshold value. The process of convergence to this solution is enhanced and a solution more likely to be found when the initial parameter values chosen are close to the 'true' values.

Some probability distributions lend themselves to the task of estimating these initial values by having functions which can be transformed into linear functions of time. For example a complementary $\log - \log$ transformation of the Weibull survival function gives,

$$\log\left(-\log\left[S\left(t\right)\right]\right) = \frac{1}{b}\log t - \frac{a}{b}$$

which is a linear function of log-transformed time, $\log t$. Hence survival data given this transformation will be approximately linear when plotted against $\log t$ if the data follow the Weibull distribution.

Linear regression can then be used to estimate values for $a$ and $b$. Such estimates should be treated as approximate as it is unlikely transformed data satisfy conditions for linear regression, e.g., concerning the distribution and variance of residuals. Furthermore the number of points over which the regression is performed is determined by the number of sampling intervals, rather than the numbers of individuals involved. Parametric non-linear regression models specified according to the log-likelihood equations described in the main text avoid these problems, as well as explicitly taking into account censored data.

Initial parameter estimates describing background mortality can be made directly from the data of uninfected hosts, as this is the only source of mortality assumed to be acting. In contrast the observed pattern of mortality for infected hosts is assumed to arise as the product of surviving background mortality and mortality due to infection. There is no reason to assume these independent and mutually exclusive sources of mortality will follow the same pattern over time. The pattern of mortality due to infection can be identified by calculating the relative survival of infected hosts and transforming these data to test for linear relationships over time. A useful exception for the need to calculate relative survival is when the observed cumulative survival of uninfected hosts and that observed for infected hosts are both roughly linear when complementary $\log - \log$ transformed and plotted against $\log\left(time\right)$. In this case the pattern of mortality due to infection can be assumed to follow the Weibull distribution as the product of two Weibull distributions also has a Weibull distribution.

An alternative to applying linear regressions to transformed data is to estimate parameter values by non-linear regression, for example, by fitting survival functions to observed survival data. Such estimates should also be treated as approximate as they respond to change in survival over time, rather than the actual frequencies of individuals dying or remaining alive in each sampling interval.

### S06 Analysis of Blanford et al. data (i)

This example analysed a subset of the data from the study by Blanford et al (1). The data came from the third experimental block and concern the survival of adult female mosquitoes exposed to the isolate *Bb06* of the fungal pathogen *Beauveria bassiana* and those in the matching control treatment. In each case, data from four replicate populations were pooled together.

Most of the females in the infected treatment died within the 14 days of the experiment, whereas roughly half of those in the uninfected cages died (Fig S6.1a). Thus by the end of the experiment the relative survival of females in the infected treatment was roughly twice their observed survival (Fig S6.1b).

The pathogen's virulence was estimated as increasing over time (Fig S6.1b), based on the dynamics of change in the relative survival of infected hosts, $h_{INF}(t) = -S'_{REL}(t)/S_{REL}(t)$.

Daily data on the frequencies of individuals dying or remaining alive in each treatment allowed the pathogen's virulence to be estimated each day ($\pm 95\% c.i.$) based on differences in the observed rates of mortality in the infected and uninfected treatments, $h_{INF}(t) = h_{OBS.INF}(t) - h_{BCK}(t)$. These estimates indicated daily rates of mortality due to infection became significantly greater than zero in the second week of the experiment, although not consistently so (Fig S6.1c).

The observed survival in the infected and uninfected treatments was roughly linear when given a complementary log − log transformation and plotted against log-transformed time (Fig S6.1d). This suggested both the background mortality and mortality due to infection were suitable for description by the Weibull distribution.

Linear regression of these transformed data provided initial estimates for the location and scale parameters for each source of mortality (Table S6.1).

Table S6.1: Location ($a$) and scale ($b$) parameters for background mortality and mortality due to infection as estimated by linear regression (LR) and maximum likelihood (ML)

| Parameter | $a$ | | $b$ | |
|---|---|---|---|---|
| | mean | 95% c.i. | mean | 95% c.i. |
| Background mortality | | | | |
| LR | 3.343 | 3.054-3.757 | 0.792 | 0.662-0.987 |
| ML | 2.845 | 2.723-2.999 | 0.483 | 0.407-0.579 |
| Mortality due to infection | | | | |
| LR | 2.508 | 2.365-2.684 | 0.493 | 0.428-0.582 |
| ML | 2.581 | 2.525-2.640 | 0.183 | 0.122-0.251 |

The log-likelihood model was parameterised with Weibull functions describing background mortality and mortality due to infection and an analysis performed using the linear regression estimates as initial values.

The parameter values obtained by maximum likelihood estimation differed from those obtained by linear regression, with only the 95% confidence intervals for the location parameter describing mortality due to infection overlapping for the two approaches (Table S6.1).

With four parameters estimated and a loss function of 646.6, the AICc for the maximum likelihood model was 1301. When the model was re-parameterised with the values estimated by linear regression the loss function rose to 690.8, giving an AICc of 1390. This difference in AICc (+88) indicates the maximum likelihood approach provided a much better description of the data than that achieved by linear regression of transformed cumulative survival data.

In numerical terms the hazard function describing the pathogen's virulence was,

$$h_{INF}(t) = \frac{1}{0.183t} \exp\left(\frac{\log t - 2.581}{0.183}\right)$$

As, $0 < b_{INF} < 0.5$, this function describes the rate of mortality due to infection as increasing monotonically over time at an accelerating rate (Fig S6.1e).

In contrast to estimates at individual points in time, the delta method estimated the lower 95% confidence interval as consistently greater than zero in the second week of the experiment (Fig S6.1e).

(1) doi:10.1186/1475-2875-11-365

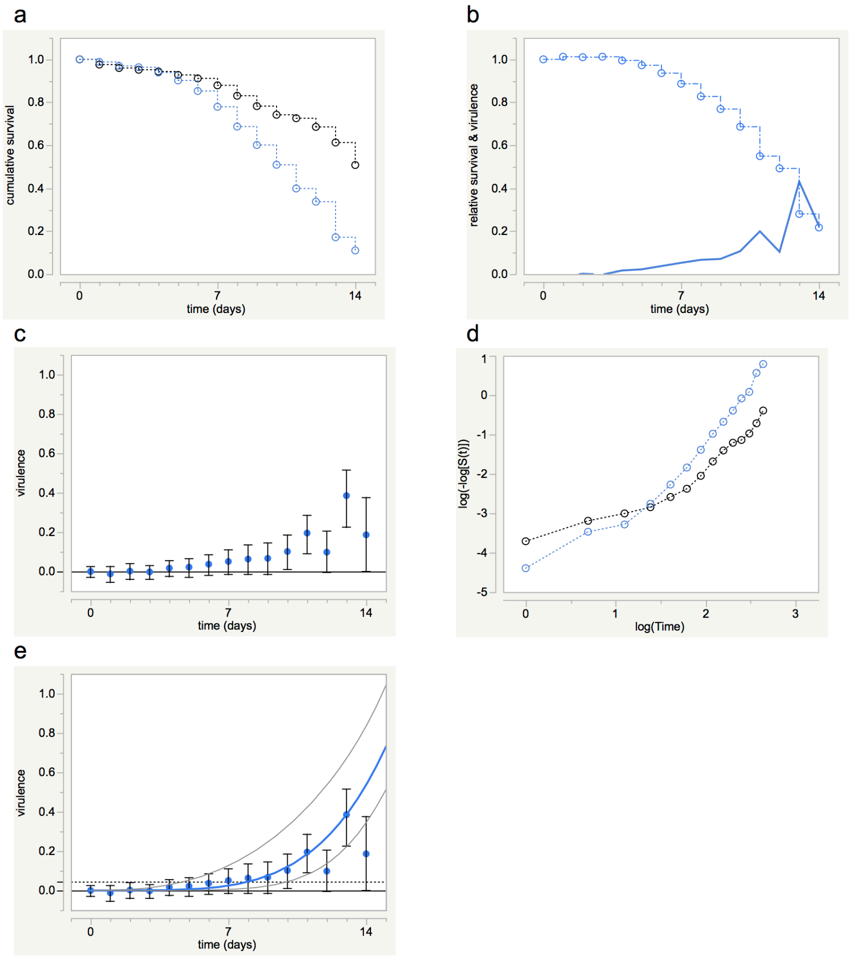


Figure S6.1: Patterns of survival and estimates of virulence for mosquitoes with fungal infections. (a) Survival of uninfected (black) and infected (blue) mosquitoes pooled across replicate cages, (b) Relative survival of infected mosquitoes (symbols/dot-dash line) and virulence estimated from dynamics of change in relative survival, (c) Virulence as estimated at individual points in time ($\pm 95\%\, c.i.$), (d) Complementary log-log transformed cumulative survival data plotted against $\log(time)$; uninfected mosquitoes (black), infected mosquitoes (blue), (e) Virulence as estimated by maximum likelihood; solid blue line ($\pm 95\%\, c.i.$, grey lines estimated by delta method), symbols ($\pm 95\%\, c.i.$) as in (c). Dashed horizontal line is estimate for virulence assuming mortality rates remain constant over time (see main text)

### S07 Estimating average longevity in *R*

The expected or average longevity of hosts in a particular population is equal to the area under the population's survival curve. This can be estimated by integrating the survival function used to describe the population.

For example, if the location and scale parameters for an uninfected population of hosts equal *a1* = 3.080 and *b1* = 0.473, the average longevity of uninfected hosts can be estimated in *R* (1) as;

```
a1 = 3.080
b1 = 0.473

Suninf <- function(t,a1,b1){
  z1 <- (log(t)-a1)/b1
  S1 <- exp(-exp(z1))
  }

est.average.longevity <- integrate(Suninf,0,Inf,a1,b1)
est.average.longevity
```

Giving,

```
19.27051 with absolute error < 5.5e-06
```

If the location and scale parameters describing mortality due to infection in a matching population of infected hosts equals, *a2* = 2.527 and *b2* = 0.198, respectively, the expected longevity of infected hosts can be calculated as,

```
a1 = 3.080
b1 = 0.473
a2 = 2.527
b2 = 0.198

Sobsinf <- function(t,a1,b1,a2,b2){
  z1 <- (log(t)-a1)/b1
  S1 <- exp(-exp(z1))
  z2 <- (log(t)-a2)/b2
  S2 <- exp(-exp(z2))
  av.long <- S1*S2
  }
est.average.longevity <- integrate(Sobsinf,0,Inf,a1,b1,a2,b2)
est.average.longevity
```

Giving

```
10.48757 with absolute error < 0.00027
```

(1) https://www.R-project.org/

### S08 Analysis of the Lorenz & Koella data

The data in this example are from the study by Lorenz & Koella (1) and are freely available at the Dryad digital repository (2). Two-day old larvae of *An. gambiae* were exposed to spores of the microsporidian parasite *Vavraia culicis* at doses of 0, 5, 10, 20, 40, 80 or 160 (x 1000) spores larva$^{-1}$. Larvae were reared in individual vials on diets of high or low food availability. As adults they remained in their vials of origin and were provided with sugar-water. Adult longevity was recorded daily until all individuals died. Only data from female mosquitoes are analysed here. Individuals exposed to infection that died more than 15 days old and harboured no spores were excluded, assumed uninfected. In total the survival of 256 individuals was analysed. Each dose treatment was involved in the following analyses, however for clarity only data from the control, lowest and highest dose treatments are presented in the accompanying figure.

Larval food availability had little effect on adult longevity, whereas being infected reduced survival and tended to do in dose-dependent manner (Fig S8.1a). Complementary log-log transformed relative survival of infected females was approximately linear when plotted against log-time (Fig S8.1b), indicating the Weibull distribution was suitable for describing mortality due to infection.

Complementary log-log transformed cumulative survival of uninfected females in the low food treatment was approximately linear when plotted against log-time, but this was not the case for females in the high food treatment (Fig S8.1c). In particular there was excess mortality at early times relative to that expected for data following the Weibull distribution. This excess was due to two females dying at 2.0 and 2.5 days post-emergence in the high food treatment; the other 16 females in the treatment all survived a minimum of 19 days. From this it might be tempting to exclude the first two females as outsiders, particularly as the data for the remaining females fitted well with the Weibull distribution. However some females died within five days of emergence in the infected treatments. These deaths could have been due to infection, but they could also have been due to background mortality. Thus excluding the two uninfected females that died early would risk underestimating the background rate of mortality. The Gumbel (or Smallest Extreme Value; SEV) distribution is suited to describing patterns of events where there are a few rare or infrequent events at early points in time.

The log-likelihood model for analysing relative survival was thus parameterised with Gumbel functions describing background mortality and Weibull functions for mortality due to infection.

Table S8.1: Summary statistics for the different likelihood models tested

| model | loss | parameters | AICc |
|---|---|---|---|
| 1 | 764.2 | 4 | 1536.6 |
| 2 | 760.5 | 8 | 1537.6 |
| 3 | 750.8 | 14 | 1531.3 |
| 4 | 735.8 | 28 | 1534.8 |
| 5 | 752.3 | 9 | 1523.3 |
| 6 | 753.1 | 5 | 1516.4 |

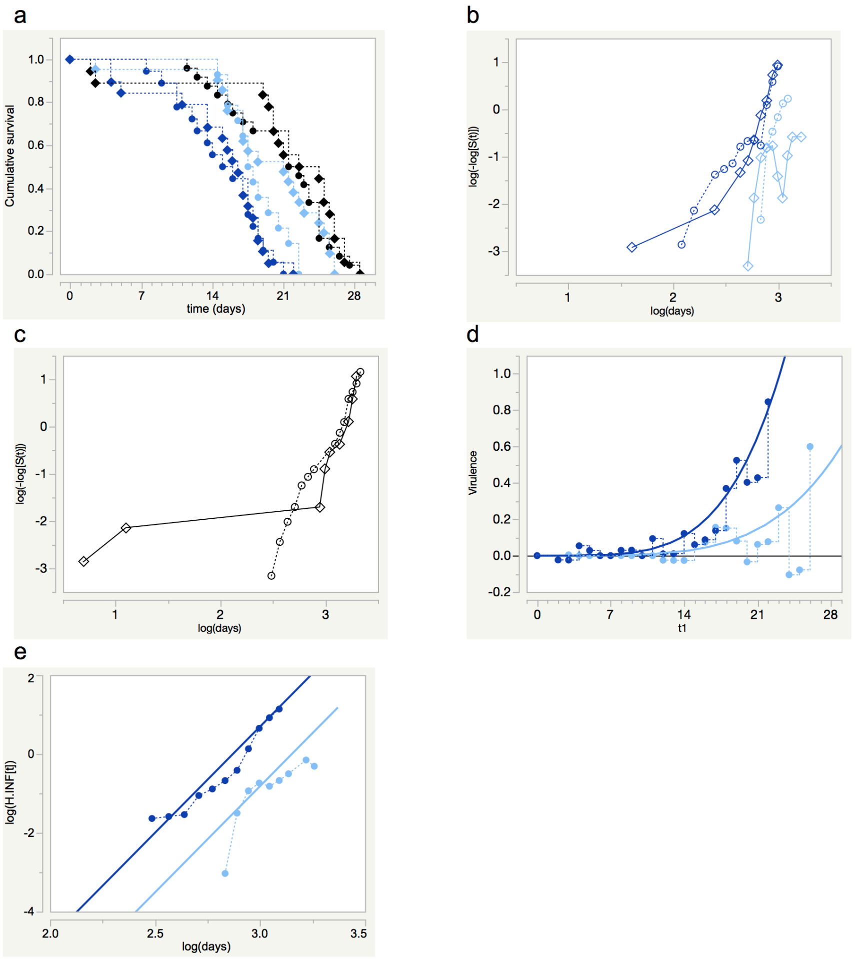


Figure S8.1: Virulence experienced by mosquitoes is proportional to dose of fungal infection. (a) Survival in control (black), lowest dose (light blue) and highest dose (dark blue) treatments. Low and high food treatments in circles and diamonds, respectively, (b) Complementary log-log (relative survival) in lowest and highest dose treatments plotted against log(time), (c) Complementary log-log (survival) plotted against log(time) for the uninfected treatments, (d) Observed and estimated virulence in the lowest and highest dose treatments. Observed data (stepped lines, symbols) based on daily data pooled for larval food treatments. Smooth curves show maximum likelihood estimates for virulence. (e) Parallel lines showing proportional nature of estimated relationship between virulence and dose. Symbols ($\pm 95\%$ $c.i.$) show observed log-log cumulative hazard data, log-log($H_{INF}[t]$), plotted against log(time)

*Model 1.* This model estimated the location and scale parameters for the background mortality and that due to infection ($a_{BCK}$, $b_{BCK}$, $a_{INF}$, $b_{INF}$). The loss function, number of parameters estimated, and the model's AICc value are given in Table S8.1.

*Model 2.* This model allowed for the effect of larval food treatments by estimating a mean value for each parameter and the deviation from this value due to the food treatment, e.g., for the location parameter describing the background mortality

$$a_{BCK} = a_0 + match(food) \begin{cases} low & \Rightarrow a_1 \\ high & \Rightarrow -a_1 \end{cases}$$

Allowing for the effect of food did not improve the model (Table S8.1).

*Model 3.* This model estimated the effect of dose treatments by estimating an underlying value for the location and scale parameters describing mortality due to infection and the deviation from this value for each dose treatment, i.e., for $a_{INF}$,

$$a_{INF} = a_2 + match(dose \,\mathrm{x}\, 1000) \begin{cases} 5 & \Rightarrow a_{d5} \\ 10 & \Rightarrow a_{d10} \\ 20 & \Rightarrow a_{d20} \\ 40 & \Rightarrow a_{d40} \\ 80 & \Rightarrow a_{d80} \\ 160 & \Rightarrow a_{d160} \end{cases}$$

where $a_{d160} = -\left(a_{d5} + a_{d10} + a_{d20} + a_{d40} + a_{d80}\right)$.

This model was an improvement on the two previous models (Table S8.1).

*Model 4.* This model re-ran Model 3 separately for each food treatment, thus allowing for a food x dose interaction. It was calculated based on the sum of the two loss functions and the total number of parameters estimated.

Model 4 did not improve on Model 3 and so the effect of larval food availability was dropped as a factor influencing adult longevity in subsequent models. The parameters values estimated in Model 3 for the deviation from the underlying estimate of $a_{INF}$ tended to decrease as the number of spores used to infect hosts increased, whereas this was not the case for deviations from $b_{INF}$ (Fig S8.2).

*Model 5.* This improved on Model 3 by fitting only a single value of $b_{INF}$ for all six infected treatments.

*Model 6.* The model improved on Model 5 by making $a_{INF}$ a linear function of $\log(dose)$ ; $a_{INF} = a_1 \cdot \log(dose) + c_1$ where $a_1$ and $c_1$ were constants.

Model 6 was judged to be the best model; the values of the parameters estimated are given in Table S8.2.

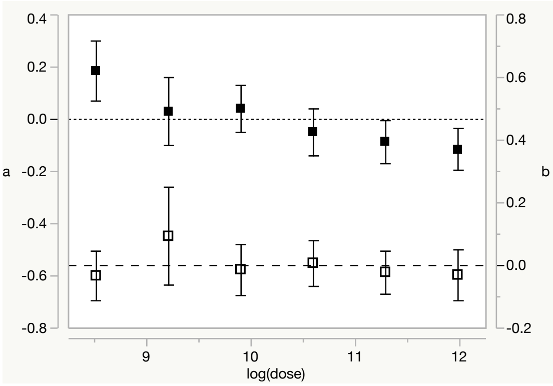


Figure S8.2: Estimated deviations from Model 3 for underlying estimates of $a_{INF}$ (left scale, closed symbols) and $b_{INF}$ (right scale, open symbols) for each dose treatment ($\pm 95\%\,c.i.$).

Table S8.2: Parameter estimates for the best likelihood model (Model 6) according to AICc

| parameter | estimate | lower 95% c.i. | upper 95% c.i. |
|---|---|---|---|
| $a_{BCK}$ | 23.182 | 21.956 | 24.565 |
| $b_{BCK}$ | 4.714 | 3.981 | 5.529 |
| $a_{INF}$ | -0.081 | -0.120 | -0.048 |
| $b_{INF}$ | 0.186 | 0.153 | 0.234 |
| $c_1$ | 3.841 | 3.494 | 4.278 |

##### Best model fitted with different probability distributions

The choice of which probability distribution(s) to use when analysing survival data determines how well the data are described. The best model above had an AICc of 1516 when the background mortality and that due to infection were described by the Gumbel and Weibull distributions, respectively. When the model was re-parameterised so the background mortality and that due to infection were described by the Weibull and Gumbel distributions, respectively, the AICc of the revised model was greater (+18), and allowing the Weibull distribution to describe both sources of mortality was worse (+25). In contrast, when the Gumbel distribution described both sources of mortality the AIC was similar to that of the best model (AICc 1518), indicating the Gumbel distribution could reasonably have been chosen to describe both sources of mortality. Had this been the case, the pathogen's virulence would still have been estimated as being proportional to log(dose), with a value of 4.476 for the ratio of virulence between highest and lowest dose treatments.

##### Proportional survival model

Could a proportional hazards model analysing the survival of infected vs. uninfected hosts have found the same results as the relative survival model? To meet the criteria for a proportional analysis, whether parametric or not, the observed rates of mortality for infected and uninfected treatments must satisfy,

$$\frac{h_{OBS.INF}(t)}{h_{BCK}(t)} = \frac{h_{BCK}(t) + h_{INF}(t)}{h_{BCK}(t)} = c$$

requiring the ratio $h_{INF}(t) / h_{BCK}(t)$ to be constant.

Thus when the background rate of mortality and that due to infection are proportional to one another, the observed rate of mortality in the infected treatments will also be proportional to the background rate of mortality. In these conditions the observed survival data are suitable for analysis with a proportional hazards model, otherwise they are not.

(1) doi: 10.1111/j.1752-4571.2011.00199.x
(2) doi: 10.5061/dryad.2s231

### S09 Accelerated failure time (AFT) model

Accelerated failure time (AFT) models satisfy the relationship that survival in one population at time $t$ equals survival in another population at time $t/c$,

$$S_A(t) = S_B(t/c)$$

where $A$ and $B$ are independent populations and $c$ is a constant scaling the passage of time in population $B$ relative to $A$. In other words, the trajectory of survival in the two populations is identical but occurs on different timescales. The hazard functions for the two populations are related as,

$$h_A(t) = (1/c)h_B(t/c)$$

which is satisfied for the Weibull distribution when the scale parameters in the two populations are equal; $b_A = b_B = b$. Hence when a Weibull survival model satisfies the conditions of a proportional hazards model it also satisfies conditions for an AFT model. This is because the two models only differ in how they describe the location parameter. For example, a complementary $\log - \log$ transformation of the Weibull survival functions $S_A\left(t\right)$ and$S_B\left(t/c\right)$ above gives,

$$\log\left[-\log S_A\left(t\right)\right] = \left(\frac{\log t - a_A}{b}\right)$$

and

$$\log\left[-\log S_B\left(t\right)\right] = \left(\frac{\log\left[t/c\right] - a_B}{b}\right) = \left(\frac{\log t - \log c - a_B}{b}\right)$$

hence

$$a_A = a_B + \log c$$

that is the two approaches differ only in describing the location parameter as a single constant or as the sum of two constants. Solving the latter for $c$ gives,

$$c = \exp(a_A - a_B)$$

hence the constant scaling the passage of time in one population relative to the other depends on the difference in the location parameters estimated for each population in a PH model.

For example, in the analysis of the Lorenz & Koella data (S8) the location parameter for mortality due to infection, $a_{INF}$, was estimated as

$$a_{INF} = 3.841 - 0.081 \cdot \log(dose)$$

where dose was the number of spores larvae were exposed to. Substituting this into the expression above for $c$ for the highest (160000 spores larva$^{-1}$) and lowest (5000 spores larva$^{-1}$) treatments gives,

$$c = \exp\left([3.841 - 0.081 \cdot \log(160000)] - [3.841 - 0.081 \cdot \log(5000)]\right) = 0.755$$

that is, survival due to infection in the highest dose treatment was equivalent to that in the lowest dose treatment when time was scaled by a factor of $t/c = t/0.755 = 1.32t$. In other words the pattern of relative survival in the highest dose treatment was equal to that in the lowest dose treatment when time in the lower dose treatment was sped up by a factor of 1.32 (Fig S9.1)

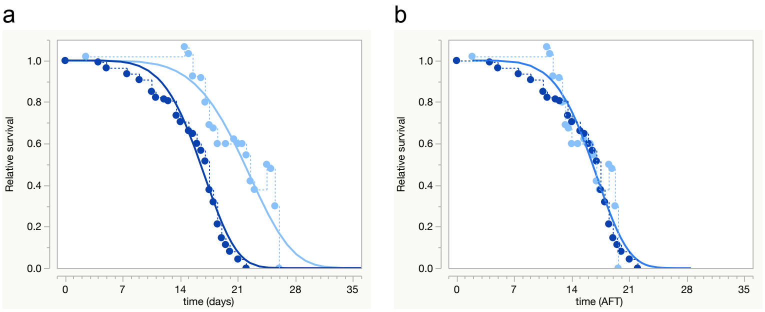


Figure S9.1: (a) Relative survival of females in the 160000 and 5000 spores larva$^{-1}$ dose treatments plotted against time (dark and light blue symbols, respectively) and (b) their relative survival when the passage of time in the low dose treatment was sped up by a factor of 1.32. Symbols and dotted lines, observed data; curves are estimated relative survival functions.

Instead of making use of the PH estimates the AFT model can be estimated directly with the log-likelihood model for estimating relative survival where the hazard function for mortality due to infection at time $t$, $h_{INF}(t)$, is

$$\begin{aligned} h_{INF}(t) &= \left(\frac{1}{c}\right)\left(\frac{1}{b_{INF}(t/c)}\right)\exp(z_{INF}) \\ &= \left(\frac{1}{b_{INF}(t)}\right)\exp(z_{INF}) \end{aligned}$$

where

$$z_{INF} = \frac{\log(t/c) - a_B}{b_B}$$

with the index $B$ defining the reference treatment against which time is scaled.

For the Lorenz & Koella data, $c$ was initially estimated as,

$$c = \exp\left( match(dose \,\text{x}\, 1000) \begin{cases} 5 & \Rightarrow a_{d5} \\ 10 & \Rightarrow a_{d10} \\ 20 & \Rightarrow a_{d20} \\ 40 & \Rightarrow a_{d40} \\ 80 & \Rightarrow a_{d80} \\ 160 & \Rightarrow a_{d160} \end{cases} - a_{d5} \right)$$

based upon $c = \exp(a_A - a_B)$, where individual location parameters were estimated for each dose treatment minus that of the reference treatment; here 5000 spores larva$^{-1}$. NB for the 5000 spores larva$^{-1}$ treatment, $c = \exp(a_{d5} - a_{d5}) = \exp(0) = 1$. Figure S9.2 shows the estimated values of $c$ for the different dose treatments.

Making $c$ a linear function of dose, $c = a_A \cdot \log(dose)$, resulted in the same parameter estimates as those of the PH model presented in the main text.

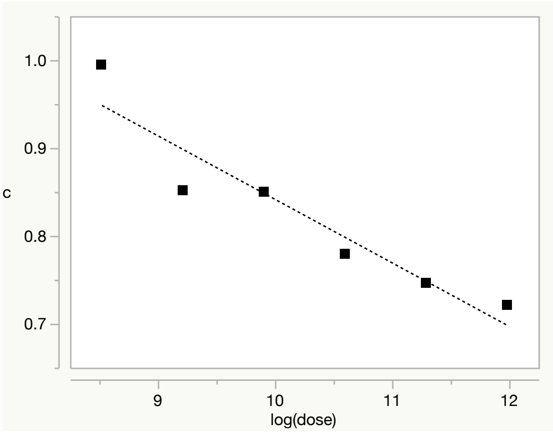


Figure S9.2: Estimated values for $c$ scaling the passage of time for mortality due to infection, taking the 5000 spores larva$^{-1}$ treatment as reference; lower values indicate less time was required for survival due to infection to equal that in the lowest dose treatment.

(1) doi: 10.5061/dryad.2s231
(2) doi: 10.1111/j.1752-4571.2011.00199.x

#### S10 Analysis of the Blanford et al. data (ii)

The data are from the study by Blanford et al. (1) and involve the fungal pathogen *Metarhizium anisopliae*. Isolates *Ma06*, *Ma07* and *Ma08* were each used to infect four replicate host populations and there were four replicate control populations unexposed to infection in the same block of the experiment. Initial populations ranged from 58 to 96 females per replicate, making a total of 1182 individuals.

Survival was reduced in the infected treatments (Fig S10.1a). The pattern of cumulative survival within replicate cages of the control treatment and when pooled together was roughly linear when given a complementary log-log transformation and plotted against log-time (Fig S10.1b), suggesting the Weibull distribution was suitable for describing background mortality. In contrast the same transformation applied to relative survival in the infected treatments was more sigmoidal or curved (Fig S10.1c), suggesting the Weibull distribution was unsuitable for describing mortality due to infection.

The flattening or leveling-off of the survival curves for infected hosts, particularly *Ma07*, indicated rates of mortality were slowing over time (Fig S10.1a). This is a pattern that can be described by the Fréchet distribution.

The log-likelihood model for analysing relative survival was parameterised using Weibull functions to describe the background mortality and Fréchet distributions to describe mortality due to infection.

Table S10.1: Estimated values for location ($a$) and scale ($b$) parameters for hazard functions describing the background rate of mortality and that due to infection for the three fungal isolates.

| Parameter | $a$ | 95% c.i. | $b$ | 95% c.i. |
|---|---|---|---|---|
| Controls | 3.503 | 3.323-3.729 | 0.686 | 0.602-0.787 |
| Isolate | | | | |
| *Ma06* | 1.872 | 1.802-1.945 | 0.514 | 0.458-0.584 |
| *Ma07* | 1.637 | 1.591-1.685 | 0.335 | 0.301-0.374 |
| *Ma08* | 2.424 | 2.262-2.630 | 0.917 | 0.764-1.123 |

The initial model estimated $a_{BCK}$ and $b_{BCK}$ for the common background mortality experienced by the mosquitoes in each treatment and separate values of $a_{INF}$ and $b_{INF}$ for each isolate (Table S10.1). As estimates of $a_{INF}$ and $b_{INF}$ for each isolate were non-overlapping the initial model was accepted as the best model.

(1) doi:10.1186/1475-2875-11-365

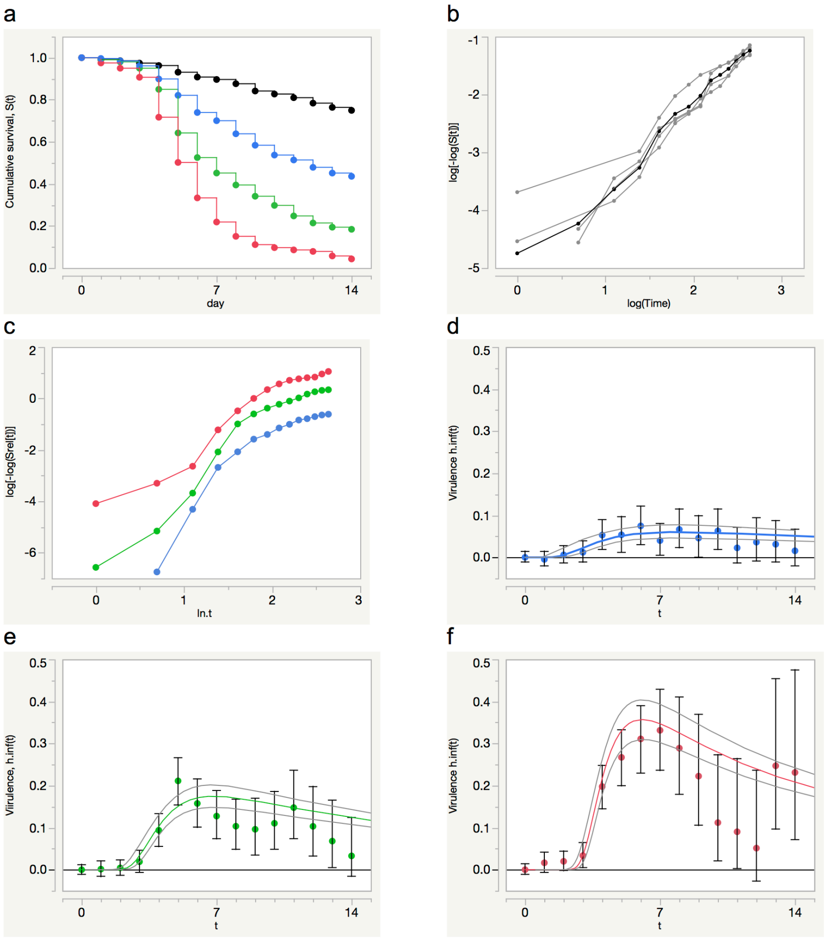


Figure S10.1: Non-proportional virulence of different fungal isolates. (a) Cumulative survival in the control treatment (black), and those exposed to isolates *Ma06* (green), *Ma07* (red), *Ma08* (blue), (b) Complementary log-log transformed cumulative survival data for the four control replicates plotted against $\log(t)$. Individual replicates shown in grey, pooled data in black, (c) Complementary log-log transformed relative survival data plotted against log(time) for the three infected treatments pooled over replicate cages, (d, e, f) Observed and estimated virulence for isolates *Ma08*, *Ma06* and *Ma07*, respectively. Observed (symbols ±95% *c.i.*), estimated (coloured curves ±95% *c.i.*grey curves)

### S11 Analysis of Parker et al. data

This supplement provides details of the models described in the analysis of the data (1) from the study by Parker et al. (2). These analyses do not take into account the effect of host genotype, the main aim is to illustrate how a unimodal pattern of virulence observed for a population of infected hosts can arise due variation in the virulence experienced by hosts within the population.

Aphid survival was reduced in a dose-dependent manner in the treatments exposed to infection (Fig S11.1a-c). The leveling-off of the survival curves in the infected treatments suggested mortality rates were slowing after peaking at around 7 days post-exposure to infection and suitable for description by the Fréchet distribution (Fig S11.1d-f). The Fréchet distribution was found to provide a better description of survival in the uninfected control treatment than the Weibull distribution (AICc 396.7 vs. 407.2). Consequently separate sets of Fréchet functions were used to parameterise the initial log-likelihood model describing background mortality and mortality due to infection.

*Model 1.* This model estimated location and scale parameters for background mortality and those describing mortality due to infection in each dose treatment, where the effect of dose treatments was estimated by their deviation from an underlying mean value;

$$x = x_0 + match(dose) \begin{cases} low \Rightarrow x_1 \\ medium \Rightarrow x_2 \\ high \Rightarrow -(x_1 + x_2) \end{cases}$$

*Model 2.* Here the effect of dose treatments on the location and scale parameters was revised as,

$$x = x_0 + match(dose) \begin{cases} low \Rightarrow x_1 \\ medium \Rightarrow 0 \\ high \Rightarrow -x_1 \end{cases}$$

thus making $x_0$ the estimate for the medium dose treatment with the effect of increasing or decreasing dose estimated as $\pm x_1$. As the low, medium and high involved doses of 8, 16 and 24 spores mm$^{-2}$ this made the location and scale parameters linear functions of $\log(dose)$.

Table S11.1: Summary statistics for likelihood models used to analyse the Parker et al. data

| model | loss | parameters | AICc |
|---|---|---|---|
| 1 | 778.60 | 8 | 1573.20 |
| 2 | 778.72 | 6 | 1569.44 |
| 3 | 756.31 | 6 | 1524.88 |
| 4 | 753.72 | 8 | 1522.95 |
| 5 | 755.12 | 5 | 1520.43 |

Model 2 was an improvement on Model 1 (Table S11.1). The estimates for the pathogen's virulence in each dose treatment are plotted in Figure S11.1d-f.

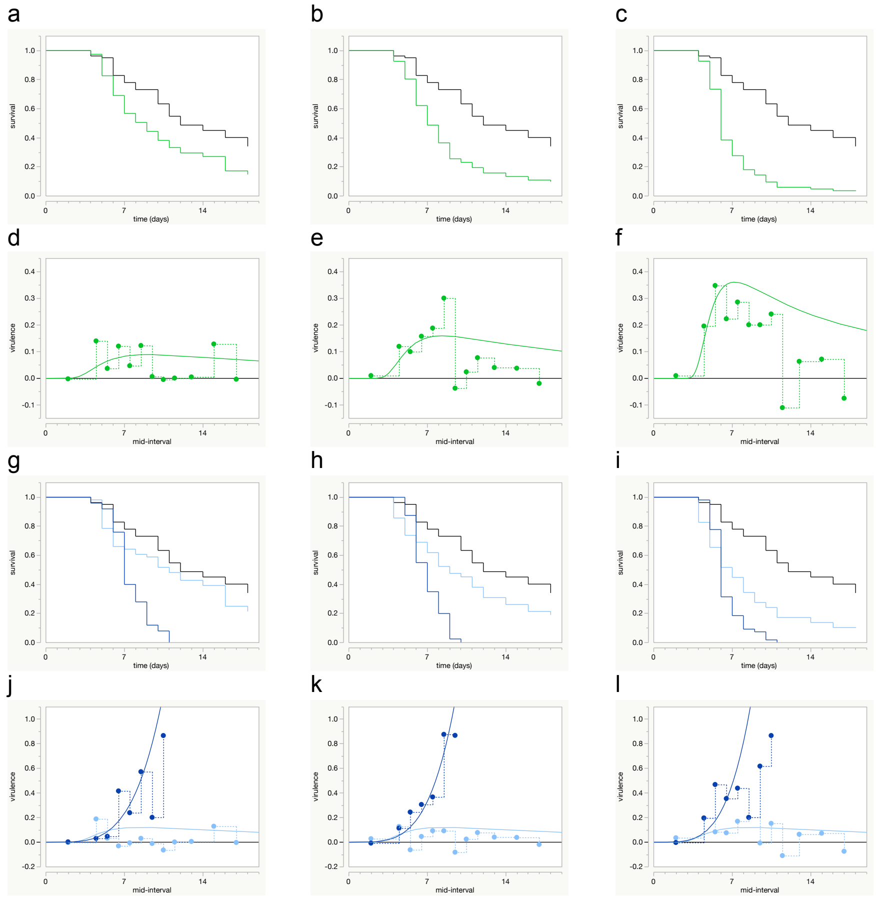


Figure S11.1: Observed unimodal patterns of virulence and underlying heterogeneity of virulence within populations. Data in the 1st, 2nd and 3rd columns are for aphids exposed to the low, medium and high dose treatments of the fungal pathogen, respectively. (a-c) Observed survival in unexposed control population (black line) and exposed population (green line). (d-f) Observed and estimated unimodal patterns of virulence at the level of the exposed population in each treatment (observed, symbols, dotted stepped line; estimated, smooth curve). (g-i) Observed survival when exposed host population classified according to sporulation status (black, unexposed controls; light blue, non-sporulating; dark blue, sporulating). (j-l) Observed and estimated virulence in sporulating and non-sporulating populations (dark blue symbols/lines, light blue symbols/lines, respectively).

*Model 3.* This model estimated location and scale parameters for individuals in the infected treatments according to whether they died during the experiment and showed signs of sporulation vs. those that did not.

$$x = x_0 + match(died\ sporulating) \begin{cases} yes \Rightarrow x_1 \\ no \Rightarrow -x_1 \end{cases}$$

These location and scale parameters were estimated using Weibull functions.

The substantial drop in the AICc values between Model 2 and Model 3 indicated classifying exposed hosts according to their sporulation status provided a much better description of the data than their dose treatment (Table S11.1).

*Model 4.* The model estimated two additional parameters where the location parameters in the sporulating and non-sporulating populations were each make linear functions of $\log(dose)$. Model 4 was a slight improvement on Model 3 (Table S11.1). Models allowing the scale parameter to vary as a linear function of $\log(dose)$ did not converge, indicating they offered no improvement to the fit of the model to the observed data.

*Model 5.* This model pooled individuals from the control treatment with the non-sporulating population of hosts exposed to infection. The sporulating population was estimated as in Model 4.

Model 5 fitted the data less well than Model 4, but it was judged a better model by AICc criteria as it estimated fewer parameters (Table S11.1). The hazard function for the pathogen's virulence in Model 5 was,

$$h_{INF}(t) = \frac{1}{b_{INF}t} \exp\left(\frac{\log t - a_{INF}}{b_{INF}}\right)$$

where,

$$a_{INF} = a_1 + \text{match}(dose) \begin{cases} low \Rightarrow a_2 \\ medium \Rightarrow 0 \\ high \Rightarrow -a_2 \end{cases}$$

the estimated parameter values are given in Table S11.2 and plotted in Figure S11.1j-l.

Table S11.2: Parameter estimates for the best likelihood model (Model 5) according to AICc

| parameter | estimate | lower ±95% *c.i.* | upper ±95% *c.i.* |
|---|---|---|---|
| $a_{BCK}$ | 2.121 | 2.044 | 2.205 |
| $b_{BCK}$ | 0.575 | 0.514 | 0.648 |
| $a_{INF}$ | | | |
| $a_1$ | 2.100 | 2.047 | 2.157 |
| $a_2$ | 0.069 | 0.009 | 0.135 |
| $b_{INF}$ | 0.197 | 0.168 | 0.232 |

These results find the hosts exposed to infection that survived until the end of the experiment or died during the experiment without showing visual signs of sporulation, had mortality rates which were no greater than those of hosts in the unexposed control population. This suggests these hosts might not have been infected, despite their exposure to infection, or that they were infected but their infections had little or no effect on their rate of mortality.

(1) doi: 10.5061/dryad.24gq7
(2) doi: 10.1111/evo.12418

### S12 Exposed-but-uninfected hosts model

Here it is assumed a 'supposedly infected' population of hosts harbours some uninfected individuals that only ever experience the same background rate of mortality as uninfected hosts in matching control treatment. These individuals may have been exposed to infection but avoided or resisted infection. Alternatively they be uninfected hosts that were never exposed to infection and accidently introduced into an infected population.

#### Infection status known

If the sub-population of exposed-but-uninfected hosts has been be identified, the log-likelihood model for estimating relative survival with right-censoring (**??** in the main text) can be used to estimate the location and scale parameters in survival functions describing background mortality and mortality due to infection,

$$\log L = \sum_{i=1}^{n} d \log \left[h_{BCK}(t_i) + g h_{INF}(t_i)\right] + \log \left[S_{BCK}(t_i)\right] + g \log \left[S_{INF}(t_i)\right]$$

where $g$ is an indicator variable taking a value of '1' for individuals exposed to infection which became infected and a value of '0' for exposed-but-uninfected individuals. $d$ is an indicator variable taking a value of '1' for individuals dying during the experiment and a value of '0' for censored individuals. $h_{BCK}(t)$ and $h_{INF}(t)$ are the hazard functions for background mortality and mortality due to infection at time $t$, respectively, while $S_{BCK}(t)$ and $S_{INF}(t)$ are the cumulative survival functions for background mortality and mortality due to infection, respectively.

This model can be run in $R$ using the code in S03.

#### Infection status unknown

The infection status of individual hosts will not always be known, but the pattern of survival observed for an infected treatment may suggest the presence of some uninfected individuals within an infected treatment, e.g., due to survival curves flattening over time. The observed patterns of survival and mortality in the 'infected' population at time $t$ can be described as,

$$\begin{aligned}
S_{OBS.INF}(t) &= p\left[S_{BCK}(t) \cdot S_{INF}(t)\right] + (1-p)\, S_{BCK}(t) \\
f_{OBS.INF}(t) &= p\left[f_{BCK}(t) \cdot S_{INF}(t) + f_{INF}(t) \cdot S_{BCK}(t)\right] + (1-p)\, f_{BCK}(t) \\
h_{OBS.INF}(t) &= f_{OBS.INF}(t) / S_{OBS.INF}(t) \\
&= \frac{p\left[f_{BCK}(t) \cdot S_{INF}(t) + f_{INF}(t) \cdot S_{BCK}(t)\right] + (1-p)\, f_{BCK}(t)}{p\left[S_{BCK}(t) \cdot S_{INF}(t)\right] + (1-p)\, S_{BCK}(t)}
\end{aligned}$$

where $p$ is a constant to be estimated, $0 \leq p \leq 1$.

##### Code for analysis in *R*

The code to run this model in $R$ (1) is given below. The model was parameterised such that functions from the Fréchet distribution are used to describe background mortality and those from the Weibull distribution for mortality due to infection.

```
exposed.but.uninfected.model <- function(t,au,bu,ai,bi,p,dataset){

  zu <- (log(t)-au)/bu
  Su <- 1-exp(-exp(-zu))
  fu <- (1/(bu*t))*exp(-zu - exp(-zu))
  hu <- fu/Su

  zi <- (log(t)-ai)/bi
  Si <- exp(-exp(zi))
  fi <- (1/(bi*t))*exp(zi - exp(zi))
  hi <- (1/(bi*t))*exp(zi)

  Sobsinf <- p*(Su*Si) + (1-p)*Su
  fobsinf <- p*(fu*Si + fi*Su) + (1-p)*fu
  hobsinf <- fobsinf/Sobsinf

  uninfected.treatment <- d*log(hu) + log(Su)
  infected.treatment <- d*log(hobsinf) + log(Sobsinf)

  logl <- -sum(fq*(ifelse(inf==0,uninfected.treatment,infected.treatment)))
  }

m01 <- mle2(
  exposed.but.uninfected.model,
  start=list(au=2.0,bu=0.5,ai=2.0,bi=0.5,p=0.5),
  data=data01)

summary(m01)
```

The model refers to four columns in the data table;
*fq* is the frequency of individuals involved,
*t* their times of death or right-censoring, and
*d* whether they died ($d$=1) or were censored ($d$=0).
*inf* is for infection status, which is used with a *ifelse* function to determine whether the expression *uninfected* or *infected* is to be evaluated.

The negative log-likelihood expression can then given starting values and evaluated by *mle2* of the *R* package *bblme* by Ben Bolker and the R Core development team (2).

##### Analysis of Parker et al. data

The model above was used to analyse the Parker et al. data (3,4). As the main effect of increasing dose was to increase the proportion of hosts experiencing virulent infections, rather than increasing the virulence of infections themselves, the data were pooled across dose treatments. The data were then analysed according to whether hosts were in the uninfected or infected treatment, where the sporulation status of individuals in the infected treatment was unknown.

Functions for the Fréchet distribution were used to estimate background mortality and those for the Weibull distribution for mortality due to infection. The loss function for the model was 790.9 giving an AICc value of 1591.9 for the 328 data and 5 parameters estimated (Table S12.1).

Table S12.1: Parameter estimates (± standard error) for the exposed-but-uninfected model for the pooled data of Parker et al.

| parameter | estimate | ± s.e. |
|---|---|---|
| au | 2.205 | 0.041 |
| bu | 0.532 | 0.030 |
| ai | 1.882 | 0.031 |
| bi | 0.167 | 0.018 |
| p | 0.480 | 0.044 |

(1) https://www.R-project.org/
(2) https://CRAN.R-project.org/package=bbmle
(3) doi:10.1111/evo.12418
(4) doi:10.5061/dryad.24gq7

#### S13 Recovery from infection model

Here recovery from infection is incorporated into a relative survival model assuming the pattern of events in an infected population at time $t$, $S_{INF.POP}(t)$, can be described as the product of three independent probability distributions,

$$S_{INF.POP}(t) = S_1(t) \cdot S_2(t) \cdot S_3(t)$$

where $S_1(t)$ is the survival function for background mortality at time $t$, $S_2(t)$ is the survival function for mortality due to infection at time $t$, and $S_3(t)$ is the survival function for the probability an infection 'survives' until time $t$, i.e., the host has not recovered at time $t$. Here the index $_{INF.POP}$, is used rather than, $_{OBS.INF}$, as recovery from infection may not be an observed event.

Differentiating the above expression with respect to time and taking the negative gives the probability density function, $f_{INF.POP}(t)$, for events occurring in the population at time $t$,

$$\begin{aligned} f_{INF.POP}(t) &= f_1(t) \cdot S_2(t) \cdot S_3(t) \\ &+ f_2(t) \cdot S_1(t) \cdot S_3(t) \\ &+ f_3(t) \cdot S_1(t) \cdot S_2(t) \end{aligned}$$

where the sum of the first two expressions gives the probability an infected host dies at time $t$, while still infected. Together they correspond with data collected for the time of death of infected hosts.

The third expression describes the probability an infected host is alive and recovers from infection at time $t$. It corresponds with data collected on the timing of recovery of infected hosts. This will not be the case if a host's recovery status is only determined after the host has died or been censored, as the data collected correspond with the time hosts recovered and subsequently survived until dying or being censored. However it is assumed recovered individuals experience the same background mortality as uninfected hosts in a matching control treatment. This pattern of mortality can be used to estimate the likelihood a recovered individual dying at time $t$, recovered at an earlier time and then survived until time $t$, when it died or was censored.

For example, in an experiment recording survival daily, the probability a recovered individual dies on the second day $(t_2)$ can be estimated as,

$$\begin{aligned} &[f_3(t_2) \cdot S_1(t_2) \cdot S_2(t_2)] \cdot h_1(t_2) + \\ &[f_3(t_1) \cdot S_1(t_1) \cdot S_2(t_1)] \cdot [S_1(t_2)/S_1(t_1)] \cdot h_1(t_2) \end{aligned}$$

where the first line gives the probability an individual recovers on the second day multiplied by the background rate of mortality on day 2. The second line gives the probability an individual recovered on the first day, survived background mortality from day 1 to day 2, $S_1(t_2)/S_1(t_1)$, and died of background mortality on day 2. Hence observed data for the times when recovered individuals die or are censored can be used to estimate the unobserved distribution of recovery times, $f_3(t)S_1(t)S_2(t)$.

When there is recovery from infection, the time when $H_{OBS.INF}(t) = 1$ and $S_{OBS.INF}(t) = 0.368$ is still potentially useful for comparative purposes. However the time when $H_{OBS.INF}(t) = 1$ is no longer when, $H_{BCK}(t)+H_{INF}(t) = 1$, or, $H_1(t) + H_2(t) = 1$, in the notation above. Instead the cumulative exposure to the risk of dying from infection needs to correct for the probability hosts recover from infection,

$$\int_0^t h_1(t) + h_2(t) \cdot S_3(t)\ dt = 1$$

which is the cumulative sum of background mortality plus that due to infection for hosts that are still infected.

Where recovery from infection occurs, and can be identified, it will also be possible to compare the numbers of infected vs. recovered individuals contributing to the overall mortality in the infected treatment when $S_{OBS.INF}(t) = 0.368$.

##### Code for analysis in *R*

The code to run this model in *R* (1) and to analyse the data using the *bbmle* package by Ben Bolker and the *R* Core development team (2) is given in a separate file (R-recovery-model.txt). A worked example and details of how to specify the data file are given further below.

##### Analysis of Parker et al. data

In this analysis of the pooled Parker et al. data (3,4), Fréchet distribution functions were used to describe background mortality and Weibull functions to describe mortality due to infection and the recovery from infection.

The loss function for the model was 937.9 giving an AICc value of 1888.1 for the 328 data and 6 parameters estimated (Table S13.1). This model fitted the pooled Parker et al. data less well than the exposed-but-uninfected model with an AICc of 1591.9 (S12).

Table S13.1: Parameter estimates (± standard error) for the recovery model for the pooled data of Parker et al.

| parameter | estimate | ± s.e. |
|---|---|---|
| a1 | 2.080 | 0.040 |
| b1 | 0.555 | 0.033 |
| a2 | 2.165 | 0.032 |
| b2 | 0.176 | 0.016 |
| a3 | 2.152 | 0.060 |
| b3 | 0.614 | 0.079 |

#### Recovered vs. exposed-but-uninfected hosts

It will not always be possible to distinguish between exposed-but-uninfected hosts and recovered hosts. However it may be possible to distinguish between populations of exposed-but-uninfected hosts and recovered hosts based on the distribution of their ages at death.

If hosts can avoid or resist becoming infected when exposed to infection, without this incurring any costs to survival, the distribution of ages at death of these exposed-but-uninfected hosts should equal that in a matching population of uninfected hosts. This will also be the case for recovered hosts if recovery from infection is instantaneous and without incurring any costs to survival. However this will not the the case if it takes time to recover from infection.

When it takes time to recover from infection, individuals are more likely to contribute towards the recovered population the later they die. Relative to a matching population of uninfected or exposed-but-infected hosts, the individuals not contributing towards the distribution of recovered individuals will be those with early ages at death, e.g., because they died of infection before recovering. This will have an effect of truncating the distribution of ages at death in the recovered population away from earlier ages at death, but not later ages at death. Consequently the mean age at death in a recovered population is expected to be later than in a matching population of uninfected or exposed-but-uninfected hosts and to have a smaller variance.

#### Recovery file details

##### Specifying the data file

The recovery model requires the data file to be specified with 10 columns named as,
*control.d,*
*control.c,*
*infected.d,*
*infected.c,*
*recovered.d,*
*recovered.c,*
*censor,*
*d,*
*t,*
*fq*

The first six columns use a combination of '1's and '0's to specify the six possible categories of hosts, as follows;
1,0,0,0,0,0 : control individuals that died during the experiment
0,1,0,0,0,0 : control individuals that were censored
0,0,1,0,0,0 : infected individuals that died while still infected
0,0,0,1,0,0 : infected individuals censored while still infected
0,0,0,0,1,0 : recovered individuals that died during experiment
0,0,0,0,0,1 : recovered individuals that were censored

The 7th column '*censor*' uses '1' to code for individuals that were censored, otherwise '0'.
The 8th column '*d*' uses '1' to code for individuals that died during the experiment, otherwise '0'; it is the complement of the '*censor*' column.
The 9th column '*t*' is for the time of the observations, e.g., in days. NB this must be an integer and there must be line for each time between $t = 1$ and the last time of sampling, $t = t_{max}$, for all 6 categories of hosts; do not include a line for $t_0$.
The 10th columnn '*fq*' codes for the frequency of individuals in each category of hosts, including '0's.

The file '*R-recovery-data-file.pdf*' provides an example of data coded in this way and to provide an example analysis. Its contents need to be saved in a file named '*R-recovery-data-file.dat*' to run the model below.

**Running the model**

(i) Change the working of directory of *R* to where the data file, '*R-recovery-data-file.dat*' is held.

(ii) Copy and paste the contents of the file '*R-recovery-model.txt*' into *R*

This will create the various functions needed for the recovery model to run and perform an analysis on the data in the file '*R-recovery-data-file.dat*' using the package '*bbmle*' by Ben Bolker (2).

The data in the file '*R-recovery-data-file.dat*' are simulated and loosely based on data from Gervasi et al. (5). Background mortality, mortality due to infection and recovery times were each simulated according to Weibull distributions. The parameter values used to generate the data were; *a1* = 2.80, *b1* = 0.50, *a2* = 2.20, *b2* = 0.35, *a3* = 2.35, *b3* = 0.35.

The recovery model assumes background mortality, mortality due to infection and recovery rates are distributed according to the Weibull distribution; this can be changed by appropriate changes to the functions; '*survival.functions*', '*calc.f3S1S2*' and '*calc.St.Sr*'. The parameter values estimated by the model should equal those in Table S13.2.

Table S13.2: Parameter estimates for recovery data

| | Estimate | Std. Error | lower 95% c.i. | upper 95% c.i. |
|---|---|---|---|---|
| a1 | 2.817413 | 0.020930 | 2.7765511 | 2.8588220 |
| b1 | 0.456838 | 0.017250 | 0.4243728 | 0.4920590 |
| a2 | 2.421926 | 0.041126 | 2.3473672 | 2.5104079 |
| b2 | 0.318360 | 0.022074 | 0.2785871 | 0.3664888 |
| a3 | 2.544519 | 0.036482 | 2.4831253 | 2.6409698 |
| b3 | 0.362418 | 0.055935 | 0.2772404 | 0.5063117 |

Figure S13.1a shows the simulated data for the observed cumulative survival in the control and infected treatments, where the latter does not take distinguish between infected or recovered hosts.

Observed and estimated distribution for times at death in the control treatment, $f_1(t)$, are shown in Figure S13.1b.

Observed and estimated distribution for times of death of infected hosts, while still infected, $f_1(t)S_2(t)S_3(t)+f_2(t)S1(t)S3(t)$, are shown in Figure S13.1c.

The recovery model estimated the unobserved probability density function for the distribution of recovery times, $f_3(t)S_1(t)S_2(t)$, based on observed times of death (or censoring) of individuals that recovered from infection (Fig. S13.1d).

(1) https://www.R-project.org/
(2) https://CRAN.R-project.org/package=bbmle
(3) doi:10.1111/evo.12418
(4) doi:10.5061/dryad.24gq7

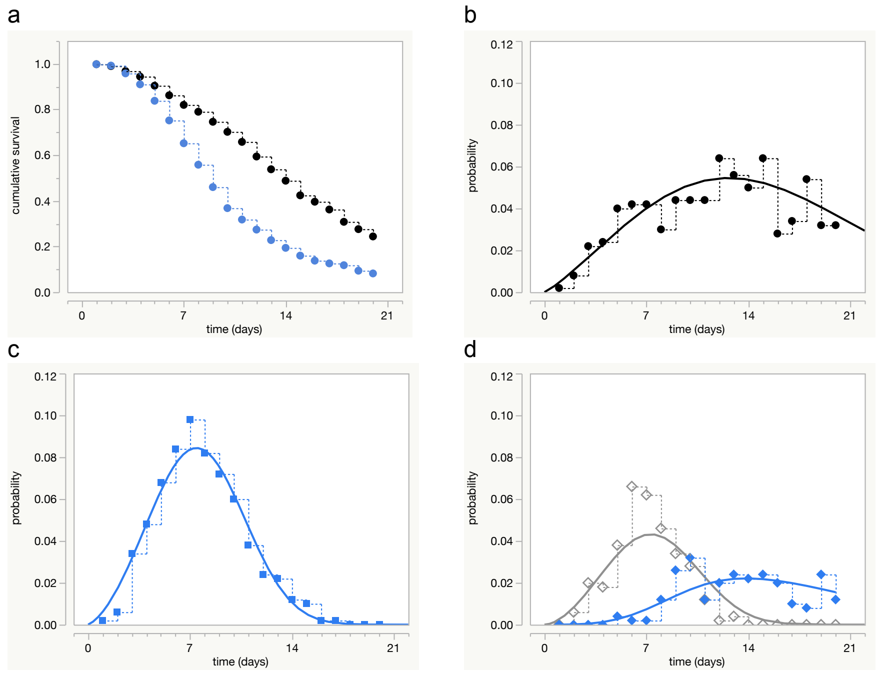


Figure S13.1: Simulated recovery data and model estimates. Simulated data (dots, stepped lines), estimated values (smooth curves). (a) cumulative survival in the control (black) and infected (blue) populations, (b) distribution of times at death in the control treatment, (c) distribution of times at death of infected individuals in the infected treatment, and (d) the unobserved distribution for the probability of recovering from infection (grey) and the observed distribution of times at death of recovered individuals (blue).

(5) doi.org/10.1098/rspb.2017.1090

### S14 Lorenz & Koella pooled data

In these analyses data (1) from the six dose treatments in the study by Lorenz & Koella (2) were pooled together to produce a single infected population in which virulence was known to be heterogeneous.

#### Virulence homogeneous

Here the infected population was modelled assuming there was no variation in virulence among infected hosts and the unimodal pattern of virulence observed at the level of the population could be described by the Fréchet distribution. The likelihood model was,

$$\log L = \sum_{i=1}^{n} \{d \log [h_{BCK}(t_i) + g h_{INF}(t_i)] + \log [S_{BCK}(t_i)] + g \log [S_{INF}(t_i)]\}$$

where the survival and hazard functions, $S_{BCK}(t)$ and $h_{BCK}(t)$, describing background mortality at time $t$ were those of the Gumbel distribution. The survival and hazard functions $S_{INF}(t)$ and $h_{INF}(t)$ describing mortality due to infection at time $t$, respectively, were those of the Fréchet distribution. $d$ is a death indicator taking values of (0,1) for individuals that were censored or died, respectively, and $g$ an indicator of infection taking values of (0,1) for uninfected and infected individuals, respectively.

Table S14.1: Estimated parameter values

| parameter | estimate | $\sim$ s.e. |
|---|---|---|
| $a_{BCK}$ | 22.694 | 0.554 |
| $b_{BCK}$ | 4.760 | 0.330 |
| $a_{INF}$ | 2.844 | 0.026 |
| $b_{INF}$ | 0.218 | 0.022 |

Table S14.1 gives the parameter values estimated by this model with their approximate standard errors. The loss function was 757.58, giving an AICc of 1523.3

#### Virulence heterogeneous

Here the infected population was modelled allowing for unobserved variation in virulence among infected hosts. This univariate frailty model assumed the unobserved variation followed the gamma distribution with a mean of 1.0 and a variance of $\theta$. The form of the likelihood model was as above, but now the hazard function describing the pattern of mortality in the infected population due to infection at time $t$, $h_{INF}(t)$ was,

$$h_{INF}(t) = \frac{h_V(t)}{1 + \theta H_V(t)}$$

where the $h_V(t)$ and $H_V(t)$ are the hazard and cumulatiave hazard functions for the underlying virulence experienced at the level of individual hosts at time $t$, respectively.

The corresponding function for the survival due to infection at time $t$ was,

$$S_{INF}(t) = [1 + \theta H_V(t)]^{-1/\theta}$$

Substituting these expressions into the likelihood model gave,

$$\log L = \sum_{i=1}^{n} \left\{ d \log \left[ h_{BCK}(t_i) + g \frac{h_v(t_i)}{1 + \theta H_V(t_i)} \right] + \log [S_{BCK}(t_i)] - g \left[ \frac{1}{\theta} \right] \log [1 + \theta H_V(t_i)] \right\}$$

The hazard and cumulative hazard functions for the underlying pattern of virulence at time $t$, $h_V(t)$ and $H_V(t)$, respectively, were those of the Weibull distribution,

$$h_V(t) = \frac{1}{b_V t} \exp(z_V) \quad \text{and} \quad H_V(t) = \exp(z_V)$$

where, $z_V = \frac{\log t - a_V}{b_V}$.

Table S14.2: Estimated parameter values

| parameter | estimate | $\sim$ s.e. |
|---|---|---|
| $a_{BCK}$ | 22.789 | 0.599 |
| $b_{BCK}$ | 4.763 | 0.340 |
| $a_V$ | 2.857 | 0.037 |
| $b_V$ | 0.090 | 0.019 |
| $\theta$ | 2.620 | 1.094 |

Table S14.2 gives the parameter values estimated by this model with their approximate standard errors. The loss function was 757.49, giving an AICc of 1525.8

(1) doi: 10.5061/dryad.2s231
(2) doi: 10.1111/j.1752-4571.2011.00199.x

### S15 Shared and correlated frailty models

Zahl (1) adapted univariate frailty models to allow for frailty effects acting on both the background rate of mortality and mortality due to infection.

#### Shared frailty

Zahl (1) cites earlier work by Hougaard (2) for this model. For a shared frailty effect the hazard function, $h(t)$, depends on the frailty effect, $\lambda$, such that,

$$h(t, \lambda) = \lambda \left[h_{BCK}(t) + h_{INF}(t)\right]$$

where $h_{BCK}(t)$ and $h_{INF}(t)$ are the background rates of mortality and mortality due to infection at time $t$, respectively. In this case the frailty effect multiplies the two rates of mortality by the same amount.

If $\lambda$ is assumed to follow the gamma distribution, with a mean of one, the hazard function for the observed rate of mortality of infected hosts at time $t$, $h_{OBS.INF}(t)$, is,

$$h_{OBS.INF}(t) = \frac{h_{BCK}(t) + h_{INF}(t)}{1 + \theta \left[H_{BCK}(t) + H_{INF}(t)\right]}$$

and their observed cumulative survival at time $t$, $S_{OBS.INF}(t)$, is

$$S_{OBS.INF}(t) = \left(1 + \theta \left[H_{BCK}(t) + H_{INF}(t)\right]\right)^{-1/\theta}$$

where the indices $_{BCK}$ and $_{INF}$ identify the background rate of mortality and that due to infection, respectively, while $\theta$ is the variance of $\lambda$ and a constant to be estimated.

The corresponding likelihood model can be written as,

$$\log L = \sum_{i=1}^{n} \left\{ d \log \left[ \frac{h_{BCK}(t_i) + g h_{INF}(t_i)}{1 + \theta \left[H_{BCK}(t_i) + g H_{INF}(t_i)\right]} \right] + \log \left[ \left(1 + \theta \left[H_{BCK}(t_i) + g H_{INF}(t_i)\right]\right)^{-1/\theta} \right] \right\}$$

where $d$ is a death indicator (0,1) for (censored, dead) individuals and $g$ an indicator of infection (0,1) for individuals in the (uninfected, infected) treatments, respectively.

Table S15.1: Parameter values estimated by the shared frailty model

| parameter | estimate | $\sim$ s.e. |
|---|---|---|
| $a_{BCK}$ | 22.896 | 0.777 |
| $b_{BCK}$ | 3.600 | 0.452 |
| $a_{INF}$ | 18.728 | 0.707 |
| $b_{INF}$ | 3.350 | 0.430 |
| $\theta$ | 0.346 | 0.160 |

This model was applied to the Lorenz & Koella data where individuals from each dose treatment were pooled into a single infected population. The Gumbel

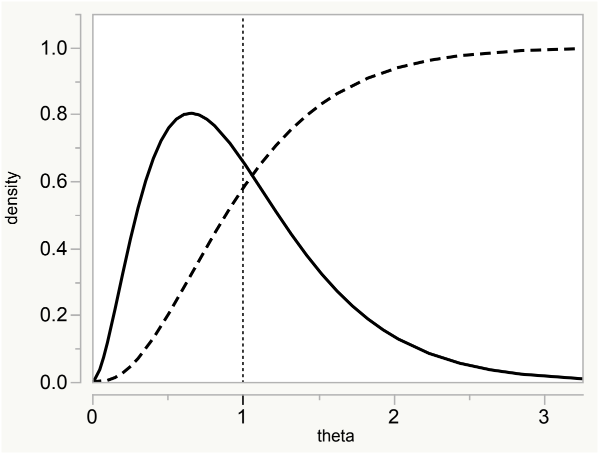


Figure S15.1: Probability density (solid line) and cumulative density (dashed line) functions for the distribution of $\lambda$ as estimated by the shared frailty model, assuming $\lambda$ follows a gamma distribution with a mean of 1.0 (vertical dotted line).

distribution was used to describe both the background mortality and that due to infection. Table S15.1 gives the parameter values estimated by this model with their approximate standard error and Figure S15.1 gives the estimated distribution of ‘frailty‘ around the value 1.0 at the beginning of the exeperiment. The loss function was 763.29, giving an AICc of 1536.8

See Zahl (1) for a discussion of the short-comings of this type of model.

#### Correlated frailty

This model allows for separate, but positively correlated, frailty effects acting on background mortality and mortality due to infection, where the strength of this correlation is a variable to be estimated.

NB the expressions for $S_{BCK}(t)$ and $S_{INF}(t)$ correct for an error in the original paper where the cumulative hazard terms were multiplied by $\sqrt{\theta_i}$, instead of $\theta_i$; $i = U, V$.

The observed rate of mortality due to infection at time $t$ in the infected population, $h_{OBS.INF}(t)$, is

$$\begin{aligned} h_{OBS.INF}(t) &= h_{BCK}(t) + h_{INF}(t) \\ &\quad - \rho\sqrt{\theta_B}\sqrt{\theta_V}\,\frac{h_{BCK}(t)\,H_V(t) + h_{INF}(t)\,H_B(t)}{1 + \theta_B H_B(t) + \theta_V H_V(t)} \end{aligned}$$

where $h_{BCK}(t)$ and $h_{INF}(t)$ are the population wide rates of mortality due to the background mortality and that due to infection at time $t$, respectively. $H_B(t)$ and $H_V(t)$ are the cumulative hazard functions for the underlying background mortality and the underlying virulence of the pathogen at time $t$, respectively; $\theta_B$ and $\theta_V$ are the variances of the unobserved variation in the background mortality and that due to infection, respectively, and constants to be estimated. $\rho$ is the strength of the positive correlation between the two frailty effects ($\rho \geq 0$).

The estimated value of $\rho$ will tend towards zero as the difference in the variance of the two frailty effects increases.

The population wide rate of background mortality at time $t$, $h_{BCK}(t)$, is

$$h_{BCK}(t) = \frac{h_B(t)}{1 + \theta_B H_B(t)}$$

where $h_B(t)$ and $H_B(t)$ are the hazard and cumulative hazard functions for the underlying rate of background mortality at time $t$, respectively. The population wide rate of mortality due to infection at time $t$, $h_{INF}(t)$, is

$$h_{INF}(t) = \frac{h_V(t)}{1 + \theta_V H_V(t)}$$

where $h_V(t)$ and $H_V(t)$ are the hazard and cumulative hazard functions for the underlying rate of mortality due to infection at time $t$, respectively.

The observed cumulative survival of infected hosts at time $t$, $S_{OBS.INF}(t)$, is given by

$$S_{OBS.INF}(t) = \left[S_{BCK}(t)^{-\theta_B} + S_{INF}(t)^{-\theta_V} - 1\right]^{-\rho/\sqrt{\theta_B}\sqrt{\theta_V}} \cdot S_{BCK}(t)^{1-\rho\sqrt{\theta_B}\sqrt{\theta_V}}\, S_{INF}(t)^{1-\rho\sqrt{\theta_V}\sqrt{\theta_B}}$$

where

$$S_{BCK}(t) = \left[1 + \theta_B H_B(t)\right]^{-1/\theta_B}$$

and

$$S_{INF}(t) = \left[1 + \theta_V H_V(t)\right]^{-1/\theta_V}$$

Table S15.2 gives the parameter values estimated by this correlated frailty model with their approximate standard errors when the lower boundary level for each parameter estimate was set to zero. The loss function was 757.59, giving an AICc of 1529.6.

Table S15.2: Parameter values estimated by correlated frailty model

| parameter | estimate | $\sim$ s.e. |
|---|---|---|
| $a_B$ | 22.789 | 0.622 |
| $b_B$ | 4.776 | 0.350 |
| $a_V$ | 16.916 | 0.642 |
| $b_V$ | 1.266 | 0.324 |
| $\theta_B$ | 0.000 | 0.027 |
| $\theta_V$ | 3.861 | 1.578 |
| $\rho$ | 0.000 | 0.281 |

The parameter values estimated by the model (Table S15.2) found there was little or no unobserved variation associated with the background mortality, $\theta_B$. In contrast, unobserved variation was estimated for the pathogen's virulence, $\theta_V$, which was known to be the case for the pooled population of infected hosts.

Correspondingly, the correlation between these two sources of frailty, $\rho$, was weak.

Setting $\rho = 0$ simplifies the model as the observed mortality rate for the infected population at time $t$, $h_{OBS.INF}(t)$, reduces to,

$$h_{OBS.INF}(t) = \frac{h_B(t)}{1 + \theta_B H_B(t)} + \frac{h_V(t)}{1 + \theta_V H_V(t)}$$

and the observed cumulative survival at time $t$, $S_{OBS.INF}(t)$, to

$$S_{OBS.INF}(t) = \left[1 + \theta_B H_B(t)\right]^{-1/\theta_B} \left[1 + \theta_V H_V(t)\right]^{-1/\theta_V}$$

When applied to the Lorenz & Koella data, this 'uncorrelated' frailty model gave the same estimates as in Table S15.2, confirming the previous estimate for there being little or no unobserved variation in the rate of background mortality.

The model allowing only for unobserved variation in mortality rates due to infection is presented in the main text.

(1) doi: 10.1002/(SICI)1097-0258(19970730)16:14¡1573::AID-SIM585¿3.0.CO;2-Q
(2) doi: 10.1093/biomet/71.1.75
