## Supplementary material for "Estimating virulence from relative survival": R recovery model and data: R-recovery-model.txt

### Recovery model ###

# This model estimates relative survival, where infected hosts can recover from infection 

# NB the dataset to be analysed needs to be specified in the function 'LL01', not in the 'mle2' command

# change the working directory of R to where the data file 'R-recovery-data-file.txt' is held
# then copy/paste the remaining text below
# this will load the data, create the various functions needed to run the model
# and analyse the data

# NB requires the package 'bbmle'


data01 <- read.table("R-recovery-data-file.txt",sep=",",header=TRUE)

head(data01)


#############################################################
######################### sub functions #####################
#############################################################

# various functions required to estimate the recovery model
# at the end of each sub-function are two lines of code that can be unchecked and run to test the individual function


##########################
### survival functions ###
##########################

# calculates various survival functions given values of location and scale parameters, a1,b1,...
# a1,b1 = location scale parameters for background mortality
# a2,b2 = location scale parameters for mortality due to infection
# a3,b3 = location scale parameters for how long infections survive

# the survival functions below are all specified according to the Weibull distribution
# these need changing to specify different probability distributions
# the functions 'calc.f3S1S2' and 'calc.St.Sr' will also need changing

calc.survival.functions <- function(a1,b1,a2,b2,a3,b3,data){

  tmax <- max(data$t)

  calc.matrix <- matrix(0,tmax,13)

  colnames(calc.matrix) <- c("t", "f1", "f2", "f3", "S1", "S2", "S3", "h1", "h2", "h3", "f1S2S3", "f2S1S3", "f3S1S2")

  for (t in 1:tmax){

  z1 <- (log(t)-a1)/b1
  z2 <- (log(t)-a2)/b2
  z3 <- (log(t)-a3)/b3

  f1 <- (1/(b1*t))*exp(z1-exp(z1))
  f2 <- (1/(b2*t))*exp(z2-exp(z2))
  f3 <- (1/(b3*t))*exp(z3-exp(z3))

  S1 <- exp(-exp(z1))
  S2 <- exp(-exp(z2))
  S3 <- exp(-exp(z3))

  h1 <- (1/(b1*t))*exp(z1)
  h2 <- (1/(b2*t))*exp(z2)
  h3 <- (1/(b3*t))*exp(z3)

  f1S2S3 <- f1*S2*S3
  f2S1S3 <- f2*S1*S3
  f3S1S2 <- f3*S1*S2

  calc.matrix[t,1] <- t
  calc.matrix[t,2] <- f1
  calc.matrix[t,3] <- f2
  calc.matrix[t,4] <- f3
  calc.matrix[t,5] <- S1
  calc.matrix[t,6] <- S2
  calc.matrix[t,7] <- S3
  calc.matrix[t,8] <- h1
  calc.matrix[t,9] <- h2
  calc.matrix[t,10] <- h3
  calc.matrix[t,11] <- f1S2S3
  calc.matrix[t,12] <- f2S1S3
  calc.matrix[t,13] <- f3S1S2
  }
 calc.matrix
}

# uncheck to test function
# test.calc.survival.functions <- calc.survival.functions(2.6,0.6,2.5,0.5,2.4,0.4,data01)
# test.calc.survival.functions

##############################
### end survival functions ###
##############################

###################
### calc f3S1S2 ###
###################

# calculate probabilities of recovery while host still alive at different times

calc.f3S1S2 <- function(a1,b1,a2,b2,a3,b3,data){

tmax <- max(data$t)

matrix01 <- matrix(0,tmax,tmax)
matrix02 <- matrix(0,tmax,tmax)

  for (t in 1:tmax){
    for (u in 1:t){
      matrix01[t,u] <- u

      z1u <- (log(u)-a1)/b1
      z2u <- (log(u)-a2)/b2
      z3u <- (log(u)-a3)/b3

      S1u <- exp(-exp(z1u))
      S2u <- exp(-exp(z2u))

      f3u <- (1/(b3*u))*exp(z3u-exp(z3u))

      matrix02[t,u] <- f3u*S1u*S2u
    }
  }
matrix02
}

# uncheck to test function
# test.f3S1S2 <- calc.f3S1S2(2.6,0.6,2.5,0.5,2.4,0.4,data01)
# test.f3S1S2

#######################
### end calc f3S1S2 ###
#######################


#############
### St.Sr ### 
#############

# calculates probability of surviving from time of recovery, S[r], to time t, S[t]: S[t]/S[r]

calc.St.Sr <- function(a1,b1,a2,b2,a3,b3,data){

tmax <- max(data$t)

matrix03 <- matrix(0,tmax,tmax)
matrix04 <- matrix(0,tmax,tmax)

  for (t in 1:tmax){
    for (u in 1:t){
      matrix03[t,u] <- t

      z1t <- (log(t)-a1)/b1
      z1u <- (log(u)-a1)/b1

      S1t <- exp(-exp(z1t))
      S1u <- exp(-exp(z1u))

      matrix04[t,u] <- S1t/S1u
     }
  }
matrix04
}

# uncheck to test function
# test.St.Sr <- calc.St.Sr(2.6,0.6,2.5,0.5,2.4,0.4,data01)
#test.St.Sr

#################
### end St.Sr ###
#################


###########################
### calc f3S1S2.(St/Sr) ###
###########################

# calculates individuals probabilities of recovering at different times and surviving to time t

calc.f3S1S2.St.Sr <- function(a1,b1,a2,b2,a3,b3,data){

  f3S1S2 <- calc.f3S1S2(a1,b1,a2,b2,a3,b3,data)
  St.Sr <- calc.St.Sr(a1,b1,a2,b2,a3,b3,data)

  f3S1S2.St.Sr <- f3S1S2*St.Sr
  }

# uncheck to test function
# test.f3S1S2.St.Sr <- calc.f3S1S2.St.Sr(2.6,0.6,2.5,0.5,2.4,0.4,data01)
# test.f3S1S2.St.Sr

###############################
### end calc f3S1S2.(St/Sr) ###
###############################


###################################
### calc row sum f3S1S2.(St/Sr) ###
###################################

# calculates sum of probabilities of recovering at different times and surviving to time t

calc.sum.f3S1S2.St.Sr <- function(a1,b1,a2,b2,a3,b3,data){

  tmax <- max(data$t)

  f3S1S2.St.Sr <- calc.f3S1S2.St.Sr(a1,b1,a2,b2,a3,b3,data)

  vector.sum.f3S1S2.St.Sr <- rowSums(f3S1S2.St.Sr)

  sum.f3S1S2.St.Sr <- matrix(vector.sum.f3S1S2.St.Sr,,1)

  }

# uncheck to test function
# test.sum.f3S1S2.St.Sr <- calc.sum.f3S1S2.St.Sr(2.6,0.6,2.5,0.5,2.4,0.4,data01)
# test.sum.f3S1S2.St.Sr

#######################################
### end calc row sum f3S1S2.(St/Sr) ###
#######################################


##########################################################################
######################## end sub functions ###############################
##########################################################################


#########################
### likelihood matrix ###
#########################

# calculates likelihoods using sub-functions above based on values a1,b1, etc...

calc.likelihood.matrix <- function(a1,b1,a2,b2,a3,b3,data){

# define maximum time
  tmax <- max(data$t)

# convert data.frame(data) to matrix
  matrix01 <- as.matrix(data01)

# calculate various survival functions (defined elsewhere) based on values a1,a2,...

  survival.functions <- calc.survival.functions(a1,b1,a2,b2,a3,b3,data)
    f3S1S2 <- calc.f3S1S2(a1,b1,a2,b2,a3,b3,data)
    St.Sr <- calc.St.Sr(a1,b1,a2,b2,a3,b3,data)
    f3S1S2.St.Sr <- calc.f3S1S2.St.Sr(a1,b1,a2,b2,a3,b3,data)
    sum.f3S1S2.St.Sr <- calc.sum.f3S1S2.St.Sr(a1,b1,a2,b2,a3,b3,data)
    h1 <- matrix(survival.functions[ ,'h1'],,1)

# create & fill matrix for likelihood calculations

  likelihood.matrix <-  matrix(0,nrow(matrix01),15)

  likelihood.matrix[ ,01] <- matrix01[ ,"t"] 
  likelihood.matrix[ ,02] <- matrix01[ ,'control.d'] * survival.functions[ ,'f1']
  likelihood.matrix[ ,03] <- matrix01[ ,'control.c'] * survival.functions[ ,'S1']
  likelihood.matrix[ ,04] <- matrix01[ ,'infected.d'] * (survival.functions[ ,'f1S2S3'] + survival.functions[ ,'f2S1S3'])
  likelihood.matrix[ ,05] <- matrix01[ ,'infected.c'] * (survival.functions[ ,'S1'] * survival.functions[ ,'S2'] * survival.functions[ ,'S3'])
  likelihood.matrix[ ,06] <- sum.f3S1S2.St.Sr * h1 # recovered, died
  likelihood.matrix[ ,07] <- matrix01[ ,'recovered.d']
  likelihood.matrix[ ,08] <- likelihood.matrix[ , 6] * likelihood.matrix[ , 7]
  likelihood.matrix[ ,09] <- sum.f3S1S2.St.Sr # recovered, censored
  likelihood.matrix[ ,10] <- matrix01[ ,'recovered.c']
  likelihood.matrix[ ,11] <- likelihood.matrix[ , 9] * likelihood.matrix[ ,10]
  likelihood.matrix[ ,12] <- likelihood.matrix[ , 2] + 
                             likelihood.matrix[ , 3] +
                             likelihood.matrix[ , 4] +
                             likelihood.matrix[ , 5] +
                             likelihood.matrix[ , 8] +
                             likelihood.matrix[ ,11]
  likelihood.matrix[ ,13] <- log(likelihood.matrix[ ,12])
  likelihood.matrix[ ,14] <- likelihood.matrix[ ,13] *  matrix01[ ,'fq'] 
  likelihood.matrix[ ,15] <- sum(likelihood.matrix[ ,14])

# return negative log-likelihood calculated in likelihood matrix
  neg.log.likelihood <- -(likelihood.matrix[1,15])

#  separately uncheck the lines below & re-run function to see the calculations for each component contributing to likelihood calculation
#  likelihood.matrix
#  matrix01
#  survival.functions

  }

# test.likelihood.matrix <- calc.likelihood.matrix(2.6,0.6,2.5,0.5,2.4,0.4,data01)
# test.likelihood.matrix

#############################
### end likelihood matrix ###
#############################

# likelihood matrix columns
# [ ,01] time
# [ ,02] for uninfected hosts that die, the probability they die at time t (f1)
# [ ,03] for uninfected hosts that get censored, the probability they survive to time t (S1)
# [ ,04] for infected hosts that die, probability of dying while still infected (f1.S2.S3 + f2.S1.S3)
# [ ,05] for infected hosts that get censored, the probability they survive to time t (S1.S2.S3)
# [ ,06] for recovered hosts that die, probability of dying at time t
# [ ,07] column with value '1' for recovered hosts that die, otherwise '0'
# [ ,08] product of columns 6 & 7 
# [ ,09] for recovered hosts that get censored, probability of surviving to time t
# [ ,10] column with value '1' for recovered hosts that are censored, otherwise '0'
# [ ,11] product of columns 9 & 10
# [ ,12] probability of events at time t collected into single column (likelihoods)
# [ ,13] log of column 12 (log-likelihoods)
# [ ,14] log-likelihoods multiplied by frequency of hosts dying/censored at time t (in matrix01)
# [ ,15] sum of log-likelihoods multiplied by frequency


############################
### likelihood function ###
############################

# function calling for estimation of likelihood

### NB the data to be analysed needs to be specified here; data=data01
### & not in the command given to 'mle2'
### otherwise the 'mle2' has trouble 'finding' the data

LL01 <- function(a1,b1,a2,b2,a3,b3,data=data01){
  calc.LL01 <- calc.likelihood.matrix(a1,b1,a2,b2,a3,b3,data)
  }

# uncheck to test function
# test.LL01 <- LL01(2.6,0.6,2.5,0.5,2.4,0.4)
# test.LL01


################################
### end likelihood function ###
################################


#####################################################################
######################## run analysis ###############################
#####################################################################

# requires package 'bbmle'

library(bbmle)

# basic analysis: starting from parameter values used to simulate the data
m01 <- mle2(
  LL01,
  start=list(a1=2.8,b1=0.5,a2=2.2,b2=0.35,a3=2.35,b3=0.35),
  )
summary(m01)


# the parameter values used to simulate the survival data were;
# a1 = 2.8, b1 = 0.5, a2 = 2.2, b2 = 0.35, a3 = 2.35, b3 = 0.35
# data were censored after controlling for mortality at t=20

############################################################################
######################## end of run analysis ###############################
############################################################################
